# The foveal visual representation of the primate superior colliculus

**DOI:** 10.1101/554121

**Authors:** Chih-Yang Chen, Klaus-Peter Hoffmann, Claudia Distler, Ziad M. Hafed

## Abstract

Processing of foveal retinal input is important not only for high quality visual scene analysis, but also for ensuring precise, albeit tiny, gaze shifts during high acuity visual tasks. The representations of foveal retinal input in primate lateral geniculate nucleus and early visual cortices have been characterized. However, how such representations translate into precise eye movements remains unclear. Here we document functional and structural properties of the foveal visual representation of midbrain superior colliculus. We show that superior colliculus, classically associated with extra-foveal spatial representations needed for gaze shifts, is highly sensitive to visual input impinging on the fovea. Superior colliculus also represents such input in an orderly and very specific manner, and it magnifies representation of foveal images in neural tissue as much as primary visual cortex does. Primate superior colliculus contains a high-fidelity visual representation, with large foveal magnification, perfectly suited for active visuomotor control and perception.

## Introduction

A defining feature of our visual system is its foveated nature. A tiny portion (∼1-2%) of our retina, fovea, contains the highest density of photoreceptors (*1*) while, at the same time, commanding large neural tissue magnification as signals traverse from retina to lateral geniculate nucleus and primary visual cortex (V1) (*2*). Such magnification confers on foveal visual input a tremendous computational advantage compared to images sampled outside fovea. As a result, we frequently move our eyes to align tiny portions of our visual environment with our fovea; fovea represents the “origin” of our visual space, and most of our visually-guided eye-movement behavior (e.g. during reading, sewing, driving) involves aligning this “origin” with what currently matters most.

Despite the importance of foveal visual processing, and despite the suitability of primate models for investigating it (*3-11*), studies of foveal visual mechanisms have become increasingly rare in the past 3-4 decades. This is primarily due to challenges associated with studying foveal vision: individual retinal ganglion cells in the foveal representation often have response fields (RF’s) that are the size of single photoreceptors (*6, 12*); thus, even tiny fixational eye movements can displace images away from RF’s. Paradoxically, a historical shift towards studying extra-foveal eccentricities in perception and cognition was simultaneously accompanied (*13*) by an assumption that fixational eye movements in between gaze shifts are irrelevant. Because we now know that this is an oversimplification (*13, 14*), there is an ever-pressing need to investigate foveal processing mechanisms, even in the presence of active fixational gaze behavior.

Such a need is especially relevant for sensory-motor areas, like midbrain superior colliculus (SC), that are at the nexus of both visual processing and eye movement generation. The SC is instrumental for generating microsaccades (*15*), which precisely re-direct gaze during foveal vision (*16*), and which also influence peripheral eccentricities (*14, 17*). However, it is not known how microscopic gaze control can be guided visually; the SC’s foveal visual representation is unexplored. In fact, countless debates have existed about whether foveal information is directly routed from retina to SC (*5, 18-23*). Nowadays, a prevailing assumption is that foveal vision is the domain of retino-geniculate pathways, whereas extra-foveal event detection for gaze shifts is the purview of SC (*24, 25*). This dichotomy gained even more traction when it was suggested that the “foveal” zone of “motor” SC is a distinct “fixation” zone preventing, rather than generating, eye movements (*26*).

Regardless of its source, primate SC does possess a visual processing machinery (*9, 23, 25, 27-31*) capable of modulating cortical areas (*32*) and influencing eye movements (*30*). Our purpose here is to characterize the physiological and structural properties of foveal vision in SC; we show that SC is as foveal a visual structure as V1.

## Results

### Primate superior colliculus is highly sensitive to foveal visual input

We recorded from 413 neurons isolated and characterized online in awake macaque monkeys N and P. We characterized RF hotspot location (the point for which maximal visual response was evoked), RF area, and other visual response properties by presenting small spots of light over an otherwise uniform gray background, and we carefully controlled for image shifts caused by incessantly occurring fixational eye movements (Materials and Methods).

We first identified SC sites containing neurons sensitive to foveal visual stimuli. Figure S1A, B (left column) shows that RF hotspots <1-2 deg in eccentricity tiled the entire foveal region contralateral to the recorded SC. This was also true for anesthetized monkeys R and F, recorded in separate experiments (Fig. S1A, B, right column; Materials and Methods). Therefore, primate SC has clear access to foveal visual input.

We examined foveal visual RF’s in the awake animals by plotting peak stimulus-evoked response as a function of stimulus location, depicted in polar coordinates (Fig. 1A, top row showing 4 example neurons). We observed small, punctate visual RF’s having high sensitivity. All 4 neurons in Fig. 1A had RF’s almost entirely contained within the rod-sparse foveola region of the retinal image (i.e. with eccentricities <0.5 deg), suggesting access to the highest acuity portion of visual input. The middle row of Fig. 1A plots the same visual RF’s using log-polar coordinates (*15*) to magnify small eccentricities, and it demonstrates that despite the proximity of all neural RF’s to gaze line of sight, the inner borders of all RF’s were very well defined. Thus, SC foveal visual RF’s sample highly specific regions of the visual image, even within foveola.

**Figure 1.**
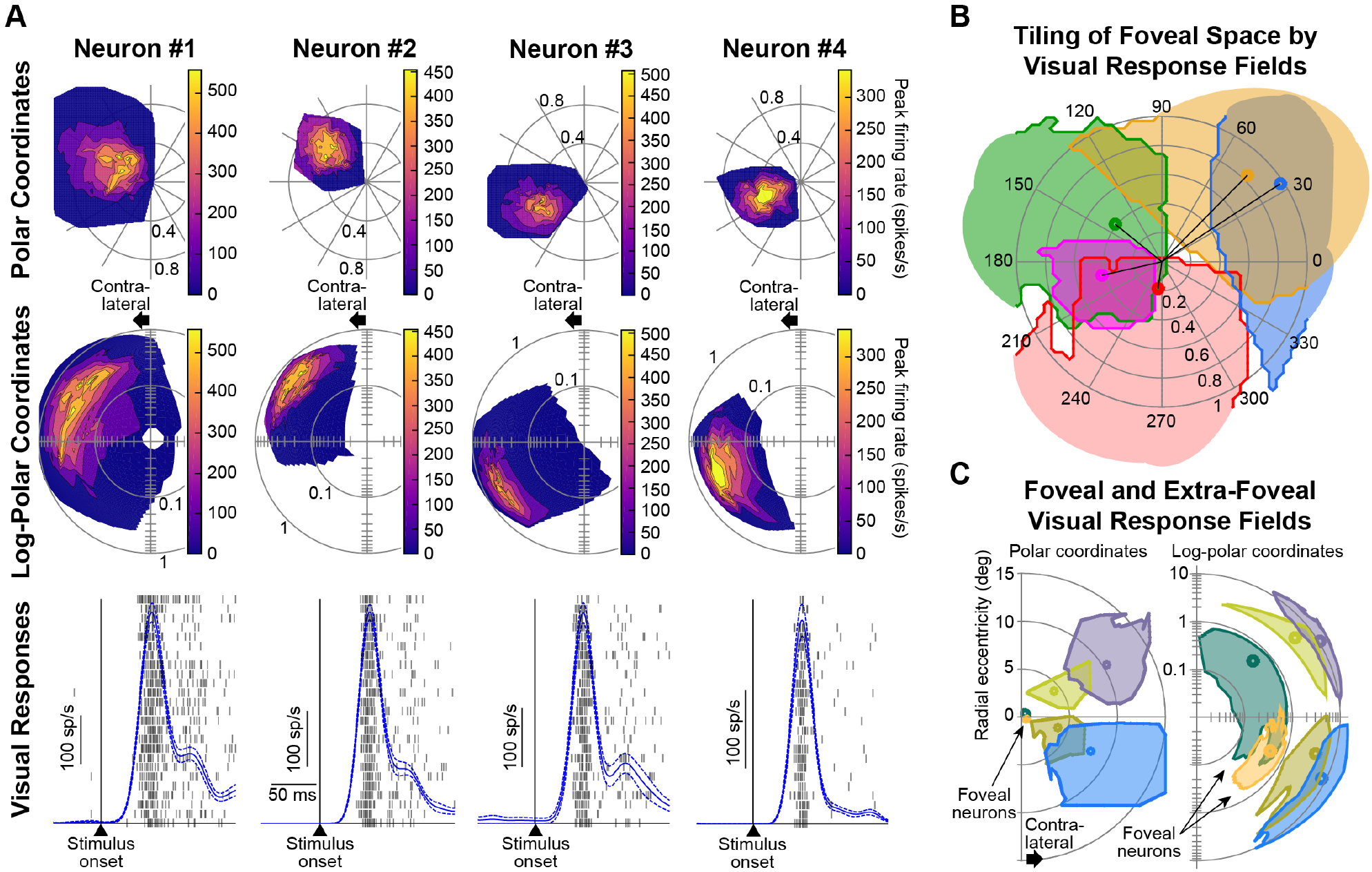
Foveal visual neurons of primate superior colliculus (SC). **(A)** Four example neurons recorded from right SC in awake monkeys. Top: visual response fields (RF’s) in polar coordinates (Materials and Methods). Middle: same RF’s using log-polar coordinates magnifying small eccentricities (*15*). Bottom: raw firing rates (blue) along with s.e.m. error bars (dashed blue) when a stimulus was presented near the neuron’s RF hotspot (Materials and Methods). Individual rasters show raw spike (action potential) times across repetitions of stimulus presentation. All neurons had RF’s almost entirely contained within the rod-sparse foveola region (i.e. <0.5 deg eccentricity; (*6-8*)). Different neurons tiled different portions of foveal space (top two rows), and all had strong sensitivity. **(B)** Across both right and left SC’s, the entire foveal space was represented. Each small circle indicates RF hotspot location from an example neuron, and solid lines indicate RF inner boundaries. **(C)** Foveal RF’s were much smaller than peripheral ones. Six example neurons with RF boundaries and hotspot locations indicated by outlines and small circles, respectively. Right: log-polar coordinates magnifying the small, but highly well-defined visual RF’s of the foveal neurons. Also see Fig. 2A.

Foveal SC neurons were also particularly light-sensitive, reaching strong peak visual responses of several hundreds of spikes/s (Fig. 1A, bottom row; each spike raster shows responses form an individual trial, and the stacked trials show responses from the 25 nearest locations to each neuron’s RF hotspot). The particular neurons shown in Fig. 1A had zero baseline activity (typical of foveal and extra-foveal SC neurons; Fig. S2A), and only responded when light impinged on a small region within fovea. Their high sensitivity was consistent across the population (Fig. S2B), and it was similar to sensitivity in more eccentric neurons (e.g. 8-10 deg in Fig. S2B). Thus, SC not only has access to even foveolar (i.e. rod-sparse) image regions, but it is also highly sensitive to them.

Strong visual sensitivity was additionally reflected in neural response latency (*29-31*): first-spike latency (Materials and Methods) was similar to that in peripheral neurons (Fig. S2C). Only in superficial SC layers, response latency decreased with increasing eccentricity within the central 2 deg (Fig. S2C, blue). Because superficial neurons, but not deeper ones, receive direct retinal input (*3-5*), this non-uniformity could reflect the sparseness of rod photoreceptors (having fast integration times) in foveola.

### Foveal superior colliculus samples visual space non-uniformly, even within foveola

The fact that foveal SC neurons have well-defined RF’s (Fig. 1A) suggests that the population as a whole can achieve full coverage of foveal image regions. We observed this not only in hotspot locations (Fig. S1), but also when we examined individual RF areas (Fig. 1B). Our recorded neurons exhibited classic hallmarks of population coding: tiling of space in terms of hotspot location as well as RF overlap, much like saccade-related activity for much larger eccentricities (*33*).

We were particularly interested in the dependence of such foveal population coding on eccentricity. It is known that RF area increases with eccentricity (Fig. 1C) (*9, 31*), like in visual cortices (*34*), but it is also assumed that spatial resolution in visual cortices is uniform within the foveal image representation (*34*). This is not what we found. There was clear dependence of RF area on eccentricity even within foveal eccentricities (Fig. 2A, left, awake animals). Moreover, this dependence followed a consistent supra-linear relationship with extra-foveal neurons (Fig. 2A, right, each dot is a neuron). Therefore, there exists a continuum of SC visual representation, from well-within the rod-sparse foveolar region up to eccentricities two orders of magnitude larger. This non-uniformity of spatial sampling even within fovea also explains why microsaccades are needed to re-align gaze (*16*): tiny microsaccade-induced image displacements equivalent to a mere handful of individual foveal photoreceptors are sufficient to bring images of fine stimuli onto SC RF’s with higher spatial resolution (smaller area) than before the eye movements.

**Figure 2.**
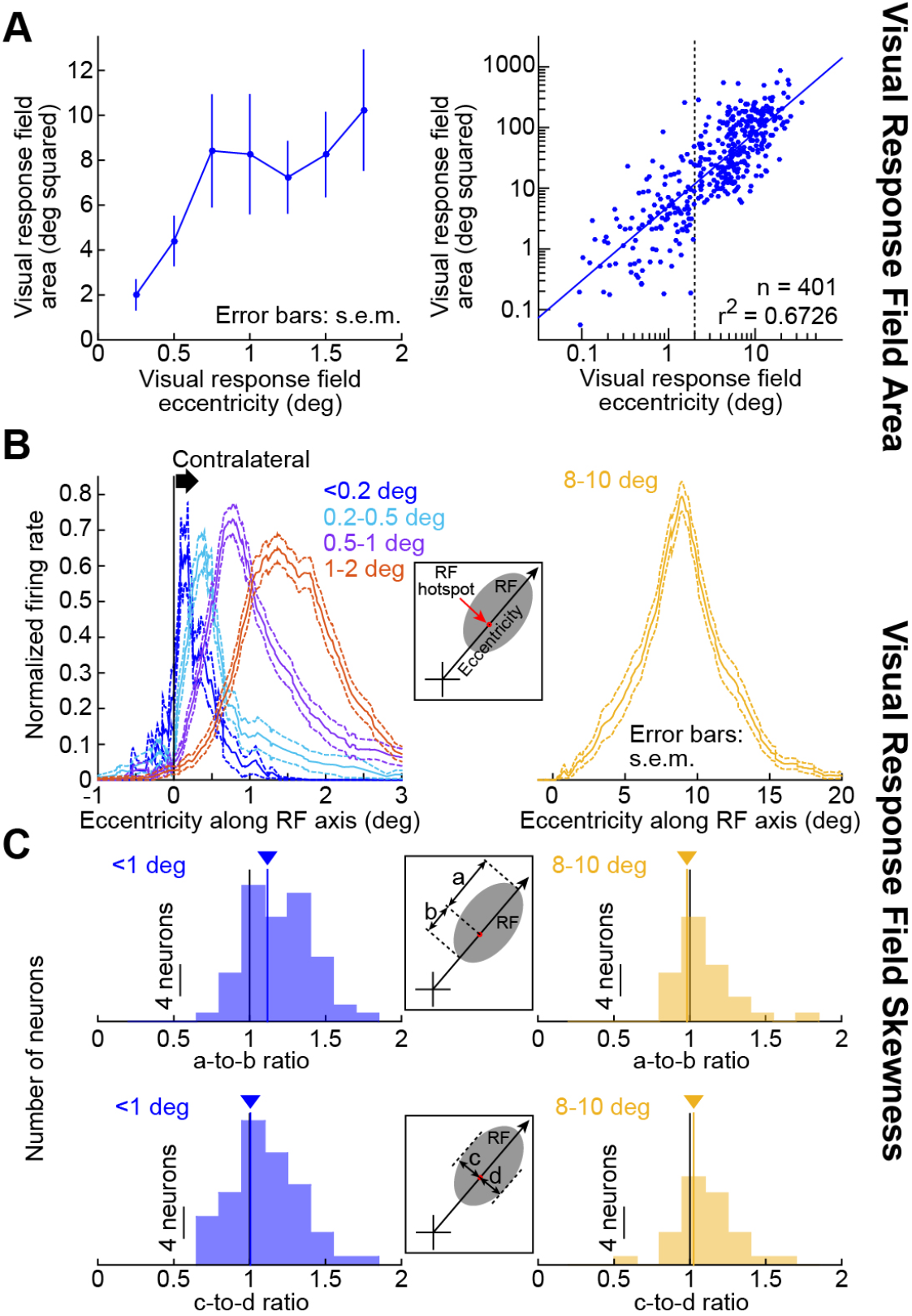
Non-uniform foveal spatial sampling by SC. **(A)** Left: visual RF area as a function of eccentricity (awake monkeys N and P). RF area depended on eccentricity (p=0.044, 1-way ANOVA, F=2.782, n=121 neurons). Right: RF areas from both foveal and eccentric neurons (dots) together. Foveal neurons (e.g. < 2 deg, dashed line) formed a continuum with peripheral ones. **(B)** Non-uniform spatial sampling even within individual RF’s. We binned neurons according to their preferred eccentricity (color-coded). For each bin, we plotted 1-dimensional normalized RF profiles along the radial axis connecting gaze center to RF hotspot (inset; Materials and Methods). Eccentric neurons (e.g. right panel) had symmetric RF’s. Foveal neurons became increasingly skewed with decreasing eccentricity; there was no activity beyond the 0-deg singularity towards the ipsilateral visual field. Each curve shows average normalized firing rate from all neurons within a given eccentricity bin, along with s.e.m. across neurons. Fig. S1 shows raw foveal hotspot locations. **(C)** Top row: RF skewness quantified as the ratio of *a*-to-*b* (inset; Materials and Methods). Foveal neurons (left) had skewed RF’s (p=3.797×10^-5^, Wilcoxon signed rank test against a median of 1); eccentric neurons (right) did not (p=0.906, Wilcoxon signed rank test against a median of 1). Bottom row: *c*-to-*d* ratios for the same neurons (i.e. the RF profiles along the direction dimension at the RF hotspot eccentricity; inset; Materials and Methods). Both foveal (p=0.504) and eccentric (p=0.367) neurons did not show significant RF skewness in the direction dimension. Solid colored lines indicate median values across the population.

We also confirmed the results of Fig. 2A in anesthetized monkeys, from which we recorded the most superficial SC visual laminae (Fig. S3; Materials and Methods). Additionally, because deeper SC layers exhibit saccade-related activity, we expected microsaccade-related activity (*15*) to emerge in the awake animals below foveal visual neurons; this was indeed the case (Fig. S4). It was also the case that microsaccade-related movement fields sampled foveal space non-uniformly like the visual RF’s (Fig. S5), but the movement fields were smaller. In all analyses, here and elsewhere, relevant statistics are shown in respective figure legends.

### Foveal superior colliculus exhibits strict spatial cut-offs in representing contralateral and ipsilateral visual eccentricity

Non-uniformity of SC spatial sampling within foveal eccentricities was not restricted to RF areas. Even individual RF’s were often strongly skewed. For example, the RF of Neuron #1 in Fig. 1A (top row) had a longer activity tail at eccentricities larger than RF hotspot location compared to smaller. We characterized such skewness by plotting RF profiles along a single dimension (eccentricity) connecting the line of sight to a given neuron’s RF hotspot (Fig. 2B, inset), and we classified neurons according to their preferred eccentricities (Fig. 2B, different colors). Extra-foveal neurons were largely symmetric along the eccentricity dimension (Fig. 2B, right, showing example neurons representing 8-10 deg). However, as RF location approached the discontinuity of “zero” eccentricity in the fovea, there was increasing RF skewness (Fig. 2B, left; compare activity beyond versus nearer than the preferred eccentricity eliciting maximal response in each curve). On the inner edges of RF’s, there was an extremely strict cutoff of neural activity, such that there was no activity on the other side of the “zero” deg discontinuity (ipsilateral to the recorded SC). This happened even if it meant strong RF skew (e.g. Fig. 2B, left, blue). Even neurons with <0.2 deg preferred eccentricity (Fig. 2B, left, blue) were strictly contralateral (any residual ipsilateral activity on the other side of “zero” was due to visual activation by the fixation spot itself, as in Fig. 5 below, or unavoidable residual eye-position calibration/correction errors). These results are consistent with a “split fovea” representation (*18, 35*).

We quantified RF skewness by measuring distance *a* from RF hotspot location to RF outer border and comparing it to distance *b* from RF hotspot location to RF inner border (Fig. 2C, top inset). Foveal SC neurons were significantly skewed (Fig. 2C, top left, *a*-to-*b* ratio>1) when compared to extra-foveal neurons (Fig. 2C, top right, *a*-to-*b* ration=1). Such skewness was restricted to the eccentricity dimension because measuring RF’s along an axis orthogonal to eccentricity (Fig. 2C, bottom inset) revealed clear symmetry whether the neurons were foveal (Fig. 2C, bottom left) or extra-foveal (Fig. 2C, bottom right). Therefore, foveal SC visual responses are strongly lateralized along the eccentricity dimension.

### Superior colliculus possesses orderly topographic representation of foveal visual space

Computational models of anatomical topography (*36*) suggest that RF skewness (Fig. 2) is a hallmark of foveal magnification. We therefore next turned to the question of how SC foveal vision is structurally represented. In anesthetized animals (Materials and Methods), we densely recorded the most superficial SC laminae (putatively, the retina-recipient neurons). We then sacrificed the animals and carefully registered our dense recordings with structural neuroanatomy (Fig. S6, Movies S1-S2, and Data Table S1; Materials and Methods). We assessed preferred eccentricities along SC surface (Fig. 3A, showing a top-down view; each gray dot is an SC electrode location, Materials and Methods). We plotted iso-eccentricity lines (color-coded) and found orderly SC topography even within the foveal representation. The orderly eccentricity organization was also continuous with extra-foveal eccentricities (Fig. 3A; thick red lines show iso-eccentricity lines in our model of SC topography, Fig. 3C). In this analysis, we were careful to obtain a view of SC similar to that used in previous estimates (*37*) of SC topography that were based on evoked saccades after electrical microstimulation of the deeper layers. This allowed us to directly compare our data, containing explicit measures of foveal topography, to a classic model (*37*) only extrapolating from extra-foveal (and saccadic) measurements. We also repeated the same analysis for iso-direction lines relative to the horizontal meridian (Fig. 3B). Finally, we confirmed that we could still find evidence of orderly topography even in our awake animals, recorded with single-electrode penetrations from session to session (Fig. S7).

**Figure 3.**
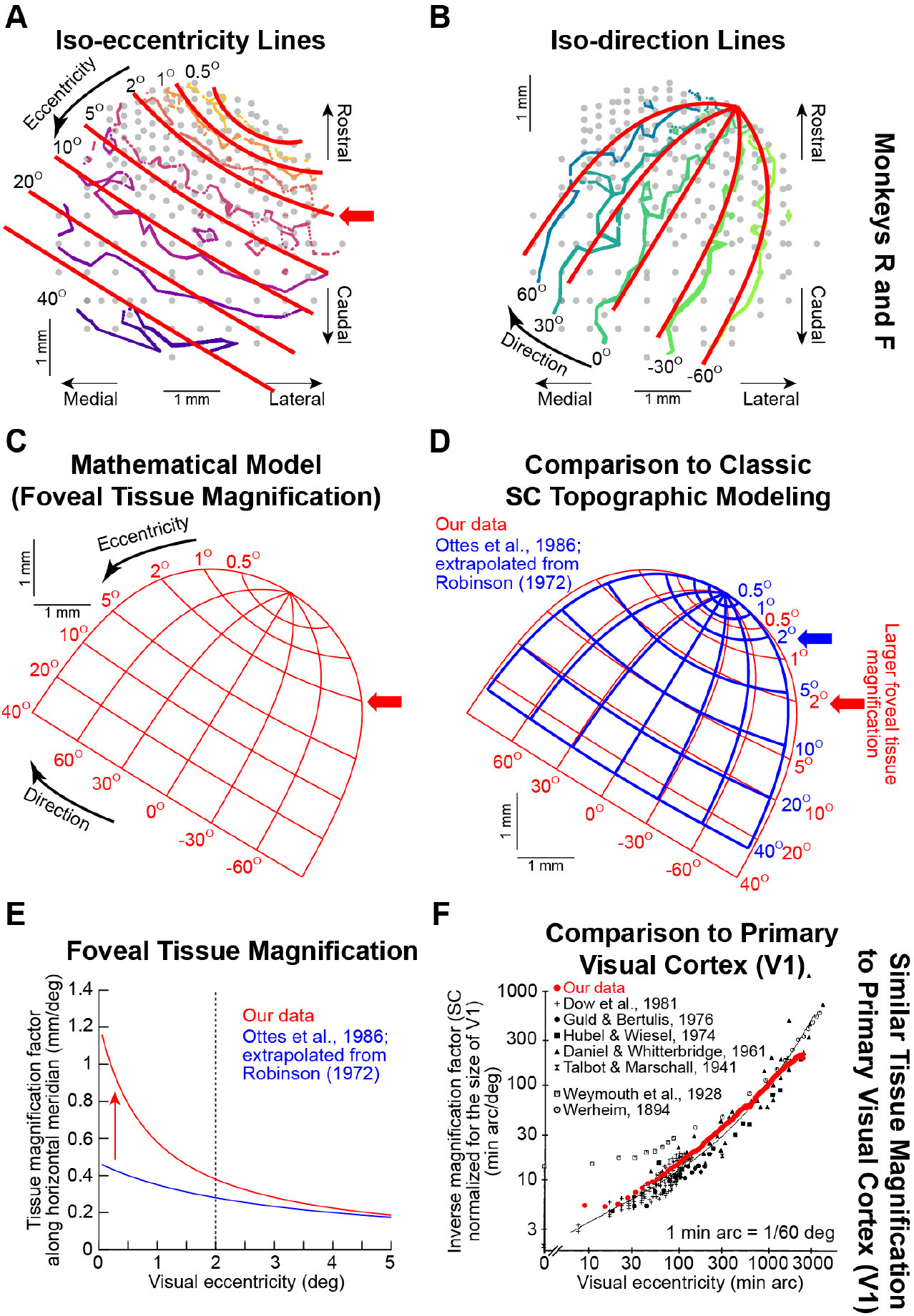
SC is as strongly foveally magnified as primary visual cortex (V1). **(A)** Using dense superficial SC sampling (anesthetized monkeys R and F), we estimated iso-eccentricity lines for right SC. Gray circles denote electrode locations (Materials and Methods; Fig. S6). We drew interpolated lines (color-coded) along which neurons preferred a single eccentricity. Foveal eccentricities were anatomically represented in an orderly manner at the rostro-lateral pole (see the 0.5, 1, and 2 deg iso-eccentricity lines). Thick red lines show iso-eccentricity lines in our mathematical model of SC surface topography (Materials and Methods; **C**). **(B)** Similar analyses for iso-direction lines. **(C)** Our SC surface topography model. The red arrow indicates 2-deg eccentricity; almost 1 mm, or approximately 1/4-1/3, of SC tissue space (along the horizontal meridian) is foveal. Data Table S1 shows SC surface topography in three-dimensional anatomical coordinates. **(D)** The universally accepted model (*37*), aligned at the same origin, assumes much smaller foveal magnification (blue map; blue arrow indicates 2-deg eccentricity). **(E)** Linear tissue magnification factor (*10, 39*) along the horizontal meridian. Our data (red) demonstrate larger than two-fold foveal magnification (vertical red arrow). **(F)** Our magnification factor curve (plotted in red as “inverse magnification factor” (*11*), in order to facilitate comparison to the other shown data) was similar to V1 foveal magnification factor (black). Note that we corrected for anatomical size differences between SC and V1 before performing this comparison to V1 (Materials and Methods). Our data (red) is superimposed on a meta-analysis adapted, with permission, from (*11*).

Therefore, foveal SC representation not only does exist (Fig. 1), but it is also organized in highly specific manners, both functionally (Fig. 2) and structurally (Fig. 3).

### Superior colliculus magnifies foveal visual representation in neural tissue as much as primary visual cortex (V1)

Our dense mappings of Fig. 3A, B, coupled with strong RF skewness (Fig. 2), convinced us that SC contains significantly larger foveal magnification than we had anticipated. We therefore fitted our data from Fig. 3A, B (color-coded lines) with a mathematical function incorporating foveal magnification (Fig. 3A, B, thick red lines), similar to past descriptions of V1 and SC (Materials and Methods). Our data fits (Fig. 3C) revealed that the 2-deg eccentricity line was located up to almost 1 mm away from the rostro-lateral “origin” of the topographic map (Fig. 3C, red arrow); this indicates that along the eccentricity dimension, up to 1/4-1/3 of the SC represents foveal eccentricities. This is larger than two-fold the magnification factor extrapolated by classic SC topographic modeling (*37*). To demonstrate this, we aligned in Fig. 3D both our data (red) as well as the original extrapolated model (*37*) (blue). We found a remarkable difference: what the original model predicts as the 5-deg eccentricity line in neural tissue is in reality only the edge of the foveal representation (2 deg).

We also explicitly calculated magnification factor. In Fig. 3E, we estimated linear foveal magnification factor along the horizontal meridian (that is, mm of neural tissue per deg of visual eccentricity) for our data (red) as well as the classic extrapolated model (*37*) (blue). We confirmed larger than two-fold increase in foveal magnification factor in fovea compared to the classic model (*37*). Note that both models converge at larger eccentricities because larger eccentricities are logarithmically compressed.

We then compared our data for SC foveal magnification to V1 foveal magnification. Because V1 is almost an order of magnitude larger than SC, we first scaled our SC measurements to equalize for overall size (Materials and Methods). We then plotted our magnification factor measurements along with an extensive meta-analysis of numerous V1 measurements (*11*) (Fig. 3F, this time plotted as “inverse magnification factor” by the authors of the meta-analysis; that is, min arc of visual eccentricity per mm of neural tissue). We found that SC foveal magnification is as good as V1 foveal magnification. In hindsight, this makes perfect sense: if SC is to visually guide highly precise microsaccadic gaze alignment (*15*), it ought to also have a high-fidelity visual representation.

An immediate functional implication of large SC foveal magnification is that small spots of light presented foveally would simultaneously activate a large population of neurons. Specifically, because each RF is spatially extended, a single stimulus is expected to activate multiple neurons, and such activation depends on RF overlap between neurons. For example, for Neurons #2 and #4 from Fig. 1, plotting their RF images in neural tissue space revealed that the area occupied in anatomical space according to our topography data in Fig. 3C was greater than five-fold the area predicted by the classic model of SC topography (Fig. 4A, compare right and left columns; the SC was rotated here to make the eccentricity dimension horizontal in the figure). This means that a small foveal spot (e.g. at 0.5 deg eccentricity) would activate an area of SC neural tissue greater than five times larger than would be classically predicted (*37*). This is significant because a well-known theoretical consequence of foveal magnification is equalization of the size of the active population across eccentricities (*37*). With the classic model, plotting RF area versus eccentricity in visual space (Fig. 4B, left) and tissue space (Fig. 4B, middle) revealed that equalization across eccentricities was only partial (the slope of the regression line in Fig. 4B, middle was >0), and foveal eccentricities were clear outliers. Only with the larger foveal magnification factor revealed by our own SC measurements could we observe complete equalization of active neural tissue (Fig. 4B, right).

**Figure 4.**
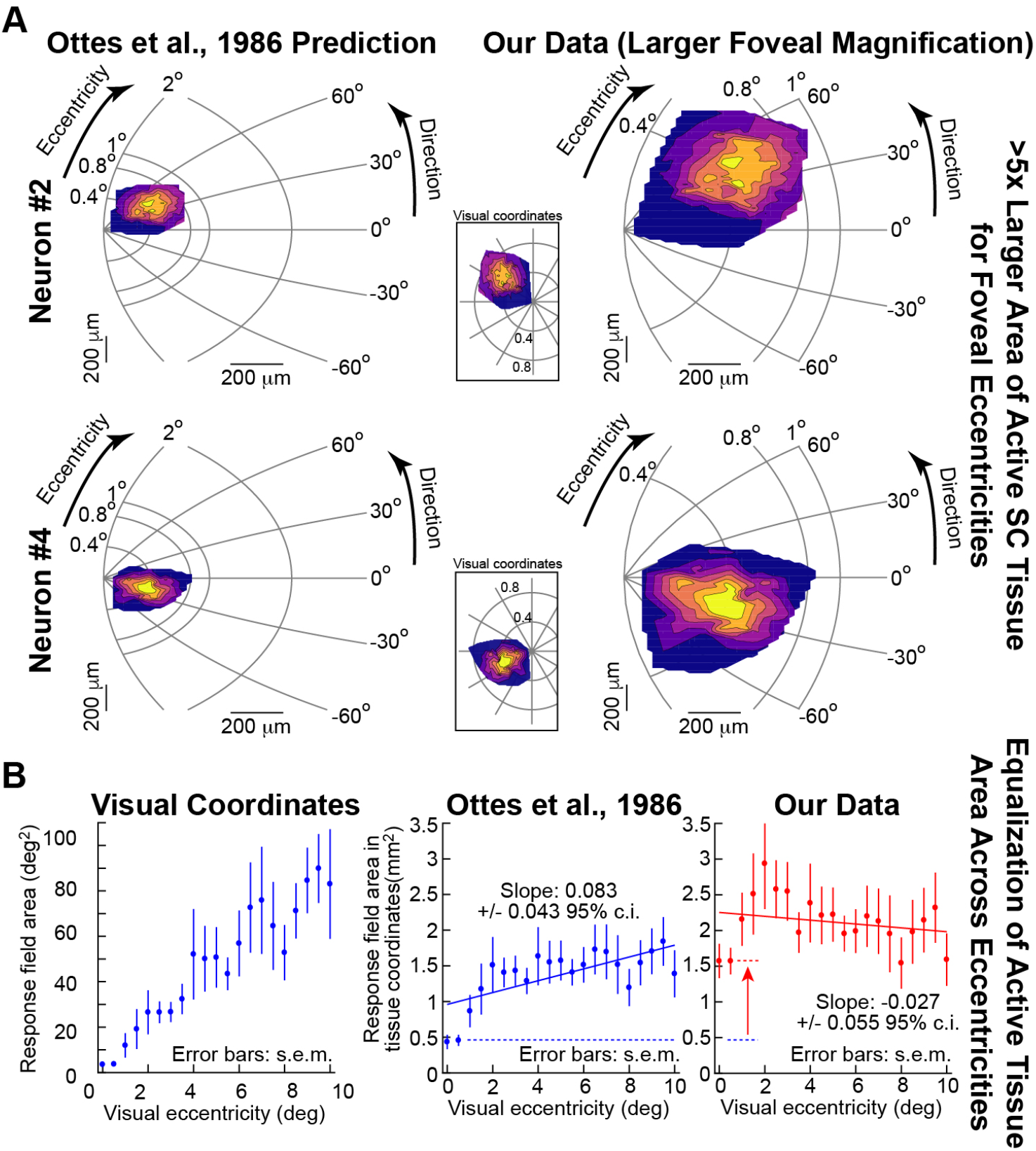
Greater than five-fold foveal tissue area magnification than previously assumed. **(A)** Each row shows the size of an example neuron’s visual RF in anatomical coordinates (left: classic SC model; right: model based on our dense sampling). This coordinate system for displaying RF’s is equivalent to estimating the SC neural tissue area activated by a stimulus (*37, 40*). The inset in each row shows each neuron’s visual RF in visual coordinates (as in Fig. 1). Our topography data (right column) demonstrate larger than five-fold active area of neural tissue for foveal visual locations than the old (*37*) estimate (left column). This is consistent with the larger linear magnification factor in Fig. 3E. Note that we rotated the SC such that the representation of the horizontal meridian (tilted in anatomy; Fig. 3) is now horizontal. **(B)** The leftmost panel shows RF areas in visual coordinates from monkey N and P (e.g. Fig. 2A). The expected increase with eccentricity should be “equalized” in SC neural tissue space due to foveal magnification (*37, 40*). However, plotting RF area in SC tissue space using the classic model (*37*) fails to result in equalization when foveal neurons are included in analysis (positive slope in middle panel; also: p=4.851×10^-8^, 1-way ANOVA as a function of eccentricity, F=4.13). This failure was also noted in (*40*). With larger foveal magnification (rightmost panel; upward red arrow), such equalization is now more successful (zero slope in rightmost panel; also: p=0.025, 1-way ANOVA as a function of eccentricity, F=1.77).

Our observation of larger SC foveal magnification not only clarifies issues related to sensory representation across the population of SC neurons (e.g. Fig. 4), but it additionally recasts interpretations of anatomical SC properties. For example, predictions on efferent projections from SC to the eye muscles (for driving saccades) (*38*) have previously relied on the classic model of SC topography. However, interpretation of the very same data would be very different based on our measurements of foveal magnification (Fig. S8). It is exactly for this reason that we provide Data Table S1 as a community resource for future studies of structure-function SC relationships.

### Fixational eye movements activate up to 1/4-1/3 of superior colliculus neural tissue by foveal stimuli

An important additional functional consequence of large foveal magnification is that small eye movements can, by virtue of moving retinal images, cause substantial variations in SC neural activity. This would happen even for tiny ocular drifts occurring in between microsaccades. Consider, for example, the scenario in Fig. 5A. A foveal RF has a well-defined inner border represented in green (consistent with Figs. 1A, B, 2B). A small ipsiversive ocular drift (away from the neuron’s RF direction; orange) would be enough to bring the image of the fixation spot into the inner edge of the RF. Thus, even when no visual stimulus is presented, the neuron might still be activated by the visual image of the fixation spot. Indeed, a fraction (22.3%) of our foveal neurons showed variable baseline activity in the absence of visual onsets (Materials and Methods). To demonstrate that this activity was visually-driven by the fixation spot itself, we identified 100-ms fixation epochs not containing any microsaccades. We then carefully assessed eye position in these epochs relative to the fixation spot (Materials and Methods). Example Neuron #5 (Fig. 5B) had an RF hotspot location of 0.3 deg. We obtained fixation epochs and sorted them based on eye position relative to the fixation spot. We also paired each epoch with simultaneously recorded spike rasters (black ticks). The neuron clearly exhibited variable baseline activity, which was strongest when eye position along the axis of the neuron’s RF hotspot (similar to Fig. 2B analysis) was ipsiversive (that is, bringing the image of the fixation spot into the RF). Thus, trial-to-trial neural variability was caused by variability of fixational eye position.

**Figure 5.**
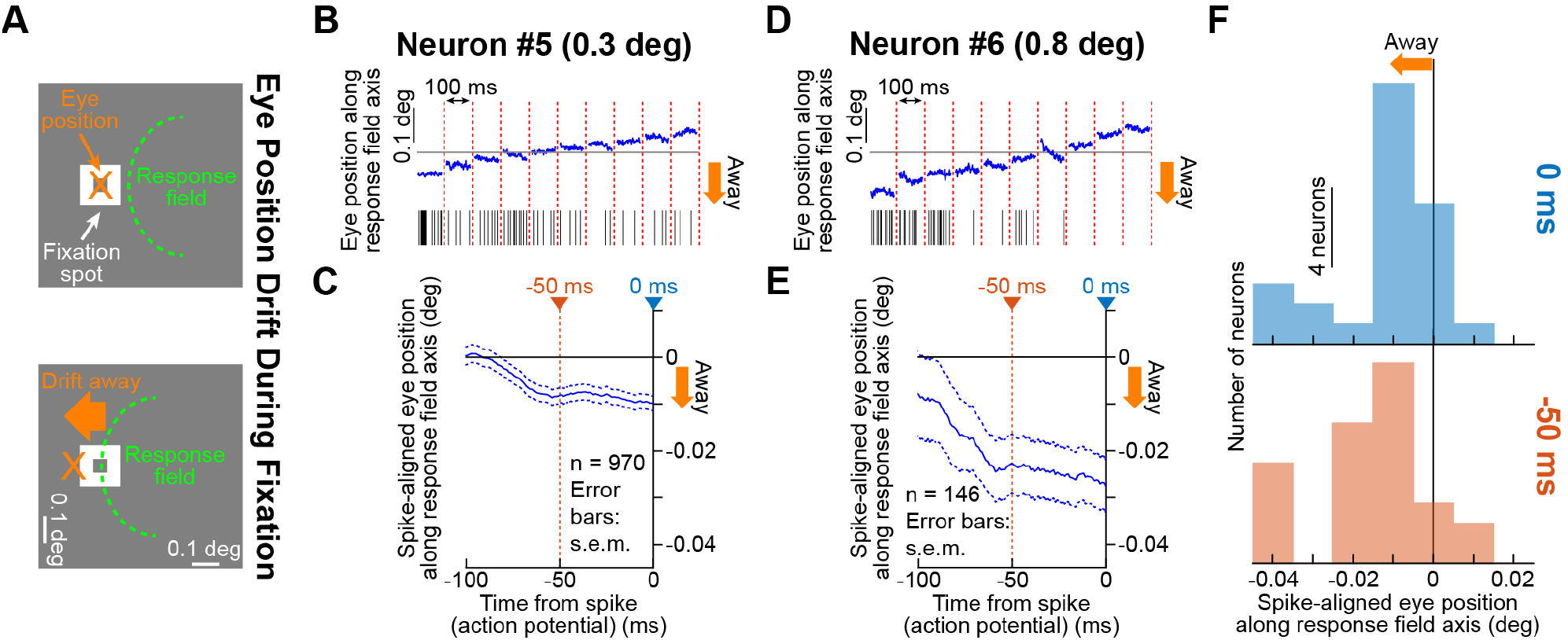
Significant impact of ocular drifts on foveal neural variability. **(A)** Top: if gaze (orange) is centered on the stationary fixation spot (white), a foveal RF (green) would not experience visual stimuli. Bottom: if gaze drifts by <0.1 deg away from the RF location, the new RF position (moving with the eye) now experiences the fixation spot as a visual stimulus. Therefore, foveal SC neuronal variability during fixation should depend on gaze. **(B)** Example neuron (with RF hotspot eccentricity at 0.3 deg) exhibiting substantial trial to trial variability in pre-stimulus activity. We analyzed eye position (blue) across trials in microsaccade-free 100-ms epochs (Materials and Methods); we sorted ten example epochs based on eye position (using the conventions of **A**). Black rasters show corresponding neural activity. The neuron was most active with eye position farthest away from the RF, consistent with **A**. **(C)** Spike-triggered average eye position during baseline fixation from the same neuron (Materials and Methods). Spikes were most likely within ∼50-100 ms after an eye position deviation away from RF location, consistent with **A**. **(D)** Similar result for a slightly more eccentric neuron; a larger ocular deviation was now needed to cause neural firing. **(E)** This was also evident in the spike-triggered average eye position. **(F)** For all neurons exhibiting baseline activity, we confirmed **B-E** by sampling the value of spike-triggered average eye position at either 0 ms (top histogram) or −50 ms (bottom histogram) relative to spike onset (colored vertical lines in **C**, **E**). Spikes were consistently preceded by “away” eye positions (for 0 ms: median = −0.0102, p=3.986×10^-5^, Wilcoxon signed rank test against 0; for −50 ms: median = −0.0127, p=4.422×10^-5^, Wilcoxon signed rank test against 0).

We further demonstrated the dependence of neural variability on eye position in this example neuron by computing spike-triggered average eye position leading up to any given spike (Fig. 5C). A spike by the neuron was more likely when eye position deviated (up to ∼50 ms earlier) away from the fixation spot (along the RF hotspot axis, consistent with the schematic of Fig. 5A). Example Neuron #6 was slightly more eccentric, and was thus activated by larger drifts in eye position away from the fixation spot (Fig. 5D, E), and measures of spike-triggered eye position were consistent across the population of neurons affording us this analysis (Fig. 5F). Therefore, because of foveal representation and magnification, significant variability of ongoing SC activity can be caused by tiny fixational gaze behavior.

If one were to now combine the results of Fig. 5 with those of Fig. 4, it would be possible to estimate, for normal fixational gaze behavior, the amount of SC neural tissue that would be activated simply when normally fixating. This amounts to almost 1/4-1/3 of the entire SC (Fig. S9). Indeed, large foveal magnification makes observations of significant influences of small eye movements on peripheral SC visual activity (*14*) reasonable.

### Microsaccades are associated with pre-movement shifts in foveal superior colliculus representations

Finally, in all of our analyses, we excluded visual onsets near the occurrence of microsaccades (Materials and Methods), because these can alter foveal visual perception (*17*). However, we had neurons in which sufficient trials with microsaccades existed, allowing us to investigate a neural basis for such foveal perceptual alteration (*17*). We mapped visual RF’s without microsaccades, and also when stimulus onset occurred <100 ms before microsaccade onset. To avoid contamination by microsaccade-related motor bursts (*15*), we only considered ipsiversive microsaccades away from RF location and not emitting an ipsiversive microsaccade-related burst. Figure 6A demonstrates results from two example foveal neurons. Consistent with perceptual alterations in (*17*), both neurons showed an ipsiversive shift in RF hotspot location (i.e. in the same direction as the microsaccade (*17*); Fig. 6A, rightward component of red arrows). Across the neurons for which we performed this analysis, there were substantial momentary shifts in RF hotspot locations if stimuli appeared before microsaccades (Fig. 6B, left). Interestingly, foveolar neurons always shifted ipsiversively (i.e. in the direction of upcoming microsaccades), whereas more eccentric neurons tended to shift outward; such a reversal with eccentricity is reminiscent of pre-microsaccadic perceptual reversals with eccentricity (*17*). Importantly, these RF shifts were not caused by differences in eye positions across the conditions (Fig. 6B, middle and right).

**Figure 6.**
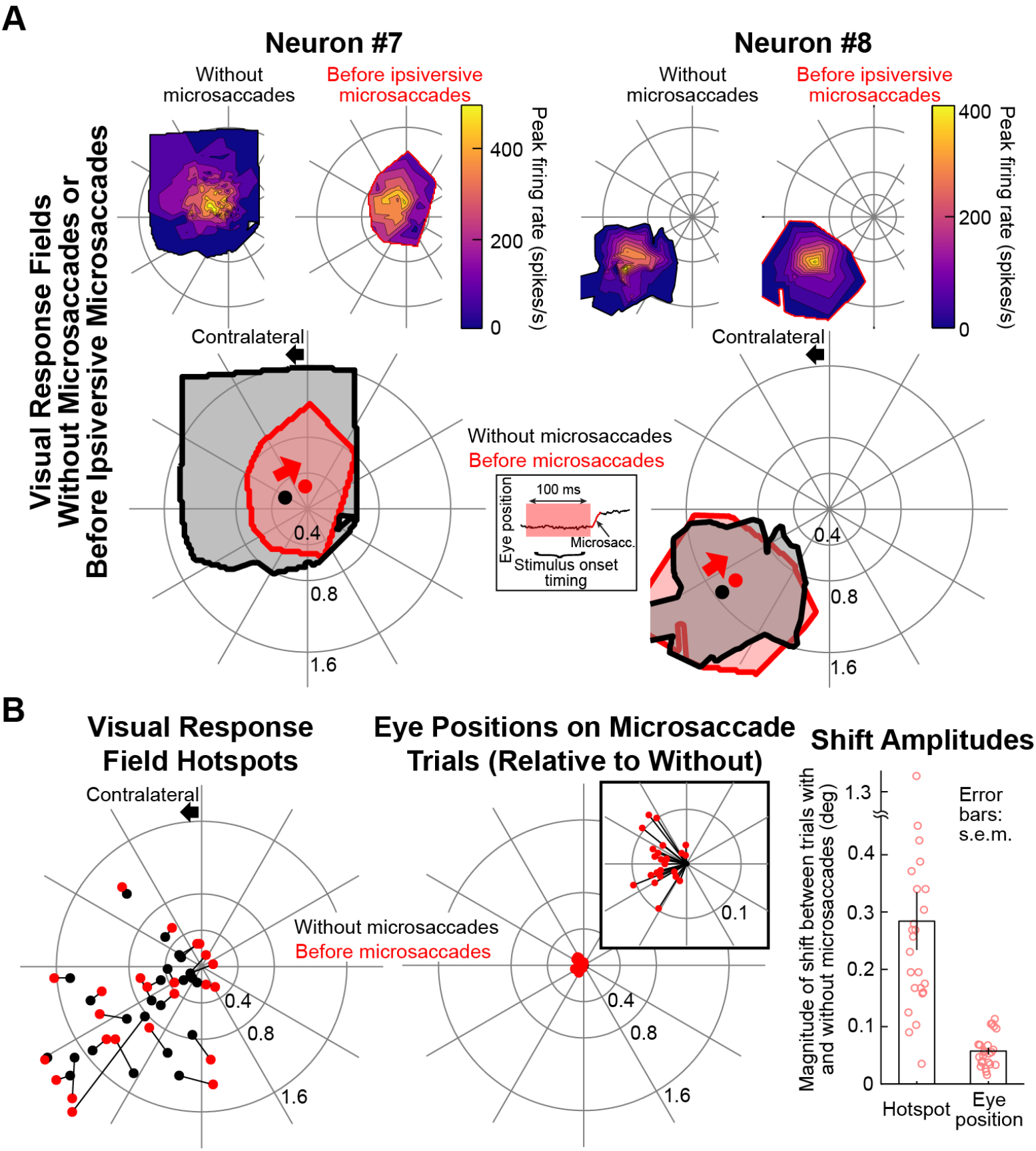
Foveal visual RF’s shift before microsaccades. **(A)** Each column shows an example neuron (#7 and #8) having sufficient trials with microsaccades to allow mapping the RF before their onset (Materials and Methods). The top left RF map in each neuron was obtained without microsaccades (like in Fig. 1). The top right map was obtained with stimuli appearing <100 ms before ipsiversive microsaccades (inset). The big panel for each neuron shows RF boundaries and centers from the two conditions. We did not compare RF boundaries between conditions because of limited numbers of microsaccades (Materials and Methods). However, substantial shifts in RF hotspot locations were clearly evident (thick red arrows). **(B)** Most foveal neurons exhibited pre-microsaccadic shifts (leftmost plot with black dots being the RF hotspot locations without microsaccades and connected red dots being the same neurons’ RF hotspot locations before microsaccades). Because microsaccades alter eye position, we also checked whether the shifts in the left panel could be explained by different eye positions at stimulus onset (middle plot). As expected (*14*), there was a small shift in eye position before ipsiversive microsaccades (inset in the middle plot), but it was significantly smaller than the RF shifts seen in the left plot. The rightmost plot summarizes the magnitude of the RF shifts (left bar) or eye position shifts (right bar) with and without microsaccades. Each circle is an individual neuron (n=24). Pre-microsaccadic RF shifts were larger than predicted by eye position shifts in the same trials (p=2.0706×10^-5^, Wilcoxon signed rank test).

Therefore, there exist not only influences of eye position on SC neural variability due to large foveal magnification (Fig. 5; Fig. S9), but microsaccades are also associated with transient changes in SC’s foveal representation (Fig. 6). This supports evidence that active fixational gaze behavior can have far-reaching impacts on neural activity and perception (*14, 17*).

## Discussion

Our results demonstrate that SC is as foveal a visual structure as V1 in terms of spatial sampling. This not only helps explain the high precision of small eye movements (*14, 16*), but it can also have implications on the role of ascending SC visual signals to extra-striate visual cortices (*32*). Such ascending signals are expected to be strongly modulated by fixational eye movements (Figs. 5, 6, S9) (*14, 17*), directly impacting how visual coding may proceed. Our topographic model of SC surface is also a useful resource for further investigations of structure-function relationships in the visual and oculomotor systems. For simplicity, we formulated the model based on the same mathematical equations as (*37*), but additional variants may be envisioned, such as including asymmetries in upper visual field representation (*31*). Anesthesia and multi-unit recording of the most superficial (retina-recipient) SC laminae in our experiments (Materials and Methods) reduced the chance of observing strong instantiations of these asymmetries in neural RF’s. Either way, the model, along with the strict lateralization that we observed in both awake and anesthetized animals (e.g. Fig. 2), also raises interesting questions on anatomical interactions between the two SC’s, for example, along the vertical meridian. In reality, we think that fixational eye movements very strongly reduce the likelihood of a purely vertical stimulus in retinotopic coordinates. Therefore, a “split” fovea representation (*18, 35*) is not as problematic as might appear at face value.

## Acknowledgments

We are grateful to H. Korbmacher and E. Brockmann for technical support. C.-Y.C. and Z.M.H. were funded by the following sources: (1) Deutsche Forschungsgemeinschaft (DFG, German Research Foundation) – Projektnummer 276693517 – SFB 1233), (2) the Hertie Institute for Clinical Brain Research (HIH), and (3) the Werner Reichardt Centre for Integrative Neuroscience (CIN). The CIN is an excellence cluster funded by the DFG (EXC 307). C.D. and K.-P.H. were funded by DFG grant Ho-450/25-1 and by ZEN grant 1.001/09/013 of the Gemeinnuetzige Hertie Foundation.

## Author contributions

C.-Y.C., C.D., K.-P.H., and Z.M.H. designed the study. C.-Y.C. and Z.M.H. conducted the awake monkey experiments. C.D. and K.-P.H. conducted the anesthetized monkey experiments. C.-Y.C., C.D., K.-P.H., and Z.M.H. analyzed the data. Z.M.H. wrote the manuscript. C.-Y.C., C.D., K.-P.H., and Z.M.H. edited the manuscript.

## Materials and Methods

### Animal preparation for the awake monkey experiments (monkeys N and P)

Macaque monkeys N and P (male, *Macaca mulatta*, aged 7 years and weighing 7-10 kg) were prepared earlier (*31, 41-43*). Briefly, we surgically attached a titanium head holder to each monkey’s skull with titanium skull screws. The head holder was used for stabilizing head position during experiments. After recovery and initial behavioral training, we implanted a scleral search coil in one eye to allow eye movement measurement using the magnetic induction technique (*44, 45*). For neural recordings from superior colliculus (SC), we implanted recording chambers, placed on the midline with chamber center being aimed at a point 1 mm posterior of and 15 mm above the interaural line (*42*). The chambers were tilted backward from vertical by 35 deg in monkey N and 38 deg in monkey P.

All experiments were approved by ethics committees at the regional governmental offices of the city of Tuebingen (Regierungspräsidium Tübingen), and they were in accordance with European Union directives on animal research, along with the German governmental implementations of these directives.

### Animal preparation for the anesthetized monkey experiments (monkeys R and F)

Two additional male macaque monkeys (monkey F, *Macaca fascicularis*, aged 13 years, weighing 6.3 kg; monkey R, *Macaca mulatta*, aged 6 years, weighing 7 kg) were used. Both monkeys had also taken part in other projects (*46, 47*). All experiments were approved by the local authorities (Regierungspräsidium Arnsberg, now LANUV) and ethics committee, and they were performed in accordance with the Deutsche Tierschutzgesetz of 7.26.2002, the European Communities Council Directive RL 2010/63/EC, and NIH guidelines for care and use of animals for experimental procedures.

The animals were pre-medicated with 0.04mg/kg atropine and initially anesthetized with 10mg/kg ketamine hydrochloride i.m. (Braun, Melsungen, Germany). After local anesthesia with 10% xylocain (Astra Zeneka, Wedel), they were intubated through the mouth, and an intravenous catheter was placed into the saphenous vein. After additional local anesthesia of their auditory canals and the skin above the planned craniotomy with bupivacain hydrochloride 0.5% (Bupivacain ^®^, DeltaSelect, Pfullingen, Germany) or prilocainhydrochloride 0.5% (Xylonest^®^, Astra Zeneca, Wedel, Germany), the animals were placed into a stereotaxic apparatus and artificially ventilated with a nitrous oxide:oxygen ratio of 3:1. Ventilation also contained 0.3-1% halothane as needed according to no change in heart rate during algetic stimuli (1% during surgery, 0.3-0.5% during recording). The skin overlying the skull was cut, and a craniotomy was performed to allow access to the SC, either vertically from above (monkey R) or at an angle of 45 deg posterior (monkey F). Heart rate, SPO_2_, blood pressure, body temperature, and end-tidal CO_2_ were monitored constantly and maintained at physiological values. Corneae were protected with contact lenses chosen with a refractometer (Rodenstock^®^) to focus the animals’ eyes on the projection screen. After surgery completion, the animals were paralyzed with alcuronium chloride (Alloferin^®^, MedaPharma, Germany).

### Histology (monkeys R and F)

Monkeys N and P are still being used for experiments. Monkeys R and F were sacrificed to reconstruct a 3-dimensional model of superficial SC surface topography (Figs. 3, 4, S6). At the end of the experiments, the animals received a lethal dose of pentobarbital (80-100mg/kg), and they were perfused through the heart with 0.9% NaCl with 0.1% procaine hydrochloride followed by paraformaldehyde-lysine-periodate containing 4% paraformaldehyde. After postfixating the tissue overnight, it was cryoprotected with 10% and 20% glycerol in 0.1 molar phosphate buffer (pH 7.4) and then flash frozen in isopentane and stored at −70°C.

Frozen sections through midbrain were cut at 40μm in the frontal plane (monkey R) or at 50μm in a plane perpendicular to SC surface (monkey F). To visualize recording sites and tracer injections (*46*), alternate series were used for Nissl stain, acetylcholine esterase (AChE) (*48*) histochemistry, cytochrome C oxidase histochemistry (*49*), myeloarchitecture, and tracer protocols (*46*). The histological sections were photographed by a photomicroscope (Zeiss Axioplan) equipped with a Canon EOS 600D camera. Brightness and contrast were adjusted with Adobe Photoshop (RRID: SciRes_000161, version 5.5).

### Behavioral tasks for the awake monkey experiments (monkeys N and P)

The animals were trained to maintain precise gaze fixation on a small spot measuring 8.5×8.5 min arc (*41*). The same animals were extensively trained on precise fixation and were used in previous studies demonstrating their high oculomotor control capabilities (*41, 43, 50-52*). The fixation spot was a white square (72 Cd/m^2^) presented over a gray background (21 Cd/m^2^) (*31, 42*). Within a given recording session, we isolated individual neurons online and characterized them using a variety of behavioral tasks. If the monkey was still interested in continuing after a neuron was recorded, we moved the recording electrode deeper into the SC and isolated additional neurons (e.g. Fig. S4). We recorded between 1 and 6 neurons per session. The behavioral tasks were chosen in part to classify neuron types and in part to characterize foveal visual neuron response properties as in the main text.

#### Memory-guided saccade task

This task was used to assist in classifying neurons as possessing a saccade-related movement response independent of visual stimulation (*53*) (see *Neuron classification* below). A fixation spot was presented for 300-1000 ms. After the monkey established gaze fixation, a second spot was flashed for ∼50 ms and disappeared. The monkey had to maintain fixation and remember the flash location. The fixation spot was then removed after 300-1100 ms; the monkey was trained to generate a saccade to the memorized flash location, and then hold gaze at that location (to within a tolerance window dependent on flash eccentricity) for ∼300-500 ms before being rewarded. Liquid reward was given if all trial epochs were successfully completed. Flash location was chosen to be at the response field (RF) hotspot of the isolated neuron, which we estimated from the other visual and saccade tasks described below. We collected >35 trials from each neuron. If the neuron elicited a saccade-related response, it was classified as being a movement-related neuron that was independent of visual stimulation (*53*) (see *Neuron classification* below).

#### Delayed visually-guided saccade task

This task was used to map visual and saccade-related RF’s. A fixation spot was presented for 300-1000 ms. After the monkey established fixation, a second spot (target) was presented. When the fixation spot was removed 500-1000 ms later, the monkey made a saccade to the target. We varied target location from trial to trial in order to map each neuron’s visual and saccade-related RF’s. For each neuron, we collected 88 +/- 34 s.d. trials. We collected data from 332 neurons in this task. Some of these neurons (mostly the peripheral ones) were used in earlier studies for other purposes (*29-31*), and they were important here as a suitable reference for comparing, in the same animals, foveal neural response properties.

#### Fixation visual RF mapping task

For the most foveal neurons, we simplified the delayed visually-guided saccade task. A fixation spot was first presented for 300 - 1000 ms. After the monkey established fixation, a second spot was then presented for 250 ms while the monkey maintained fixation. Reward was only contingent on maintaining gaze position at the fixation spot until trial end. We used this task instead of the delayed visually-guided saccade task if the neuron’s RF was too close to the fixation spot, making it difficult to instruct a delayed saccade task. Because microsaccades were inevitable in this task, this task still allowed us to explore saccade-related movement fields as well as visual RF’s, so the task was conceptually comparable to the delayed visually-guided saccade task. For each isolated neuron, we collected 122 +/- 87 s.d. trials. We collected data from 81 neurons in this task.

### Visual stimulation in the anesthetized monkey experiments (monkeys R and F)

We used small spots of light moved across the screen using a hand-held lamp or laser pointer. Single and multi-unit activity was recorded from the SC surface (upper superficial layer), and RF location and extent were determined by moving the stimulus in various directions and at various speeds across a tangent screen at a distance from the eye of 285 cm for peripheral RF’s and 570 cm for foveal RF’s. To ensure that paralysis was maintained during the experiment, a recording electrode was inserted into the visual cortex (V1) and the RF determined and checked at regular intervals during the entire recording session (also see below for RF area calculations). Horizontal RF borders were determined by moving the stimulus up and down starting outside the RF and then shifting it towards RF center. The same procedure, similar to the minimal response field method of (*54*), was also used for determining the vertical borders (this time with horizontal stimulus movements).

The exact locations of the horizontal and vertical receptive field borders in deg of visual angle with respect to the eye contralateral to the recorded SC were measured by use of a laser pointer attached to a 2D angle meter. Two calibrated rotary potentiometers aligned with the position where the eye had been during recording allowed direct reading of the azimuth and elevation values in deg.

### Neuron classification

For the anesthetized monkeys R and F, we recorded only the most superficial SC neurons. These are purely visual (*27, 55*), also receiving direct retinal projections (*3-5*). Therefore, no further classification was done. Neurons encountered at the rostro-lateral border of the SC were assigned to the nucleus of the optic tract if they responded in a direction-selective manner to temporo-nasal movement of large area random dot stimuli. Neurons at the caudal border of the SC were localized in the inferior colliculus if they responded to auditory stimuli (hand clapping, whistling, or clicks).

For the awake monkeys N and P, we classified neurons as being purely visual, visual-motor (saccade-related), or purely motor (saccade-related) (*31, 42*). We classified a neuron as being visually responsive if average neural activity 0-200 ms after target onset in the delayed visually guided-saccade task or the fixation visual RF mapping task was higher than average neural activity 0-200 ms before target onset (baseline activity) (p<0.05, paired t-test.) (*31, 42*). We also classified a neuron as being motor responsive if average neural activity 0-50 ms before saccade onset in the delayed visually-guided saccade task or the memory-guided saccade task was higher than average neural activity 100-175 ms before saccade (*31, 42*).

For some neurons, because the RF of the neuron was very close to fixation, the saccadic tasks were deemed at times to be too demanding for the monkeys. In these cases, we collected microsaccades after fixation spot onset but before the second spot onset from fixation visual RF mapping task. We classified the neurons as being motor responsive by applying the same measurement intervals above but now aligned to contraversive microsaccade onset.

In total, we analyzed data from 413 neurons from both the right and left SC (162 purely visual, 239 both visually and motor responsive, and 12 purely motor). We often combined purely visual and visual-motor neurons (401 neurons) when analyzing visual response properties, unless a specific difference was noted (e.g. Fig. S2C). We were least interested in purely motor neurons because this was not the focus in our study. Microsaccade-related responses were previously described in the SC (*15, 56*), and we used motor neurons in the present study to confirm expected vertical organization of visual, visual-motor, and purely motor neurons across the depths of the SC (*57, 58*) (e.g. Fig. S4).

### Visual and motor-related RF hotspot and area calculations

Because foveal RF’s are extremely small (e.g. Fig. 1), even the smallest fixational eye movements can smear stimulus position on the retina and therefore alter estimates of RF hotspot location and area (*59, 60*). We therefore carefully accounted for eye movement measurements in all of our experiments.

For the anesthetized animals (monkeys R and F), we used a reference electrode placed in the foveal representation of the primary visual cortex (V1) to ensure that there were no eye movements during stimulus presentation for the SC experiments; we ensured that the foveal V1 neuron was regularly stimulated (visually) after we mapped SC RF’s. Because we analyzed the entire SC surface topography estimation in a given monkey during a single session with many successive penetrations, we were severely limited in time. We therefore estimated SC RF area by listening to single or multi-unit activity, and we plotted the RF assuming a rectangular shape (e.g. inset in Fig. S3) (*54*). We demarcated the inner and outer RF locations both horizontally and vertically. Because anesthesia potentially alters RF sizes and neural response properties (*61, 62*), and because our RF area estimates under anesthesia were less quantitative than the awake monkey procedures described below, we did not perform quantitative comparisons between RF areas in the anesthetized and awake animals. However, we did confirm the same non-uniform sampling of visual space even within the foveola in both anesthetized and awake animals (e.g. Figs. 2A, S3).

For the awake monkeys (N and P), we first calibrated eye tracker measurements similar to how we calibrate them in our laboratory (and in the same animals) for our real-time gaze-contingent stimulus presentation experiments (*41, 50, 51*). We then corrected for a small technical artifact in the magnetic induction eye coil system that happened in a minority of our sessions. Specifically, for a typical ∼6-hour recording session, we sometimes observed a small, abrupt jump in eye position using the eye coil system midway through a session. We confirmed that this was a measurement artifact and not a genuine shift in the monkeys’ eye position by performing eye movement analyses (e.g. Fig. 5) and also by comparing our strong RF skewness effects (e.g. Fig. 2B) with and without artifact correction. For monkey N, we also investigated this artifact in more detail by measuring eye position using both a video-based eye tracker (EyeLink 1000, SR Research, Toronto, Canada) and the magnetic induction eye coil system simultaneously (*41*). We corrected this artifact using the following procedure. We measured average eye position 0-50 ms before stimulus onset in a given task in which a potential artifact was suspected (i.e. at the end of the initial fixation interval while the monkeys maintained stable fixation). We excluded trials with microsaccade within 0-100 ms before stimulus onset. We then used a Calinski-Harabasz criterion (*63*) on horizontal eye position, vertical eye position, and trial sequence to find an optimal clustering number, and we then ran a k-means clustering algorithm using the optimal clustering number. We validated the result by manually observing all clustering results. From among all neurons, 300 contained only 1 cluster (no artifact), and 89 contained 2 clusters. Only 24 neurons had 3 or more clusters. For the multi-cluster files, we corrected the artifact by shifting the displaced clusters to the centroid of the main eye position cloud.

With technical calibration now corrected, we were now in a position to estimate RF properties. For estimating the visual RF area, we excluded all trials containing microsaccades within +/-100 ms around stimulus onset time. We then measured peak firing rate 30-150 ms after stimulus onset. We related this visual response property to retinotopic stimulus location in order to estimate RF area in retinotopic coordinates. We obtained retinotopic stimulus location by accounting for fixational eye position variation across time and trials (*50, 51*). Specifically, we measured average eye position 0-50 ms before stimulus onset and related it to the calibrated eye position. We then shifted the true stimulus location in physical display coordinates to the retinotopic location relative to the center of gaze. For example, if the stimulus was physically presented at 15 min arc to the right of the fixation spot on the display, but the eye position at stimulus onset was shifted by 5 min arc to the left of calibrated eye position, then the true stimulus location in retinotopic coordinates was in reality at 20 min arc to the right of the center of gaze. Across all stimulus locations and peak firing rate measurements, we used Delaunay triangulation and linearly interpolated the firing rate measurements to obtain a 3-dimensional surface of visual response strength as a function of 2-dimensional stimulus location. We further applied a threshold to accept a location as being within a given RF. The threshold was that the visual response had to be >3 s.d. above baseline pre-stimulus firing rate during fixation. All locations with visual responses above this threshold were considered to be within the visual RF of a given neuron, and RF hotspot location was the location eliciting the strongest visual response. This procedure gave us a more accurate estimate of the RF area and shape, especially when the neurons were extremely close to the fovea (e.g. Fig. 2B, left panel), and we confirmed that eye position correction resulted in smaller RF’s than without correction (*59, 60*). For example, across all neurons (including eccentric ones), the average RF area decreased by 4.2 percent after correction for eye movements.

We used a similar procedure to estimate saccade-related motor RF’s. In this case, we measured average firing rate 0-50 ms before saccade onset, and we used saccadic horizontal and vertical eye position deviations as the locations against which we plotted saccade-related firing rate.

In summary analyses (e.g. Figs. 2A, S3A, S5), we binned eccentricities into 0.25 deg bin widths and used a running window of 0.5 deg step size. In Fig. 4B, for visualizing a larger eccentricity range, we used 0.5 deg bin widths and a running window of 1.5 deg step size.

### Visual RF skewness

To estimate visual RF skewness along radial retinotopic eccentricity (e.g. Fig. 2B, C), we plotted a 1-dimensional RF profile: we plotted visual response strength as a function of radial retinotopic eccentricity along the line connecting the center of gaze to the RF hotspot. To compare neurons, we normalized each neuron’s 1-dimensional RF profile along eccentricity by the neuron’s response at the peak hotspot location. We then combined neurons having a similar preferred retinotopic eccentricities (e.g. Fig. 2B). For population statistics of skewness, we fit each neuron’s 1-dimensional RF profile as a function of radial eccentricity to a beta function, and we acquired the α and β parameters of the fit. These allowed us to estimate the near and far ratios (*a*-to-*b* ratios in Fig. 2C) of the RF. We reasoned that this approach of fitting was a more unbiased method for evaluating RF skewness because the mapping could sometimes be incomplete close to the far edge of the RF; fitting takes into account the entire measurement for estimating *a*-to-*b* ratios. We also felt that the fitting gives a direct quantitative measure of RF skewness, but the skewness is also very evident in raw traces as well (Fig. 2B).

To further evaluate the above method, for the same neurons included in the current analysis, we also took another 1-dimensional RF profile, but this time as a function of direction from horizontal along the radial eccentricity of the RF hotspot (e.g. *c*-to-*d* ratio in Fig. 2C). The prediction would be that the ratio of the α and β parameters in the beta function fit would now be close to 1 because the asymmetry that we observed in RF’s was only observed along the radial dimension (Fig. 2B, C). In Fig. 2C, all neurons with preferred visual eccentricity <2 deg were analyzed, except for 6 neurons. These neurons were excluded because the RF data fit to a beta function did not result in >80% explained variance by the fit.

### Visual response sensitivity and latency

In Fig. S2B, we used the highest visual response within a neuron’s RF (i.e. at the RF hotspot) as the preferred eccentricity of each neuron. We then binned eccentricities into 0.25 deg bin widths, and we used a running window of 0.5 deg step size, to plot visual sensitivity as a function of preferred eccentricity. We combined neurons from different depths below SC surface because we did not find any significant difference in SC visual response sensitivity across depth from the SC surface. For example, comparing visual sensitivity of neurons with <1000 μm depth below SC surface to visual sensitivity of neurons with 1000-2000 μm depth resulted in a non-significant effect (Ranksum test, p=0.90848, all neurons preferring <2 deg eccentricity).

To estimate a neuron’s response latency, we first applied Poisson spike train analysis to find the first spike occurring in response to stimulus onset for each trial (*29-31, 64*). This method was particularly suitable when a neuron had an ongoing baseline firing rate on which a stimulus-evoked response rode; the method also trivially detected first-spike latency when there was no baseline rate before stimulus onset (e.g. Neurons #1-#4 in Fig. 1). To ensure that we were estimating first-spike latency for optimal stimuli across neurons, we estimated each neuron’s response latency by averaging first-spike latency only from trials in which the stimulus was near the RF hotspot (i.e. evoking a response larger than half of the peak visual response within a neuron’s RF; typically >22 trials). After we estimated each neuron’s response latency for the optimal stimulus location, we plotted the response latency across neural preferred eccentricity using 0.25 deg bin widths and a running window step size of 1 deg. We separated superficial (<1000 μm below SC surface) and deeper (1000-2000 μm below SC surface) neurons in this analysis because we observed a difference in how response latency depended on eccentricity within foveal space (< 2 deg) as a function of neural depth below SC surface (Fig. S2C).

### Neural modulations during ocular drift fixational eye movements

To demonstrate that tiny fixational eye movements can have a significant influence on neural variability in the foveal representation, even when no stimuli are presented on the display except for a fixation spot, we identified neurons having a baseline firing rate during fixation (Fig. 5). We further only considered trials in which there were no microsaccades for at least 100 ms before stimulus onset (i.e. at the end of the baseline fixation period). In these trials, the only stimulus on the display was the fixation spot (Fig. 5A); our goal was to investigate whether drifts of the image of the fixation spot into and out of a foveal neuron’s visual RF (because of fixational eye movements) would be sensed by the neurons.

We analyzed eye movements along the axis connecting the center of gaze to the RF hotspot location; that is, we rotated the eye position and RF data based on each neuron’s RF hotspot location such that the RF hotspot in the new coordinate reference frame was always to the right of the center of gaze (Fig. 5A). This allowed us to investigate eye position variation to or away from the RF (because after rotation, the horizontal eye position change was going to move the fixation spot in or out of the RF, the vertical eye position component was orthogonal to the RF and was less relevant for this analysis; we therefore excluded it from analysis for simplicity).

Next, we filtered our neurons to those having RF hotspot locations being <1 deg in eccentricity and also firing >50 spikes in the final 100 ms before stimulus onset. The latter criterion allowed us to have a more sensitive analysis for this particular question that we were interested in addressing. We also excluded neurons with microsaccades in the critical analysis period. This resulted in 27 neurons for this analysis.

Across trials, we sorted microsaccade-free eye positions based on whether they brought the fixation spot towards the RF border (Fig. 5B, top, “Away” eye positions) or away from it. If the neuron was sensitive to the influence of ocular drift on retinal images, then more action potentials should be triggered whenever the image of the fixation spot was brought towards the RF location (Fig. 5B, C, top). To plot spike-aligned eye position (Fig. 5B, C, bottom), we took every neural action potential (spike) in the final 100 ms of fixation before stimulus onset, and we plotted the average eye position (along the RF axis) leading up to this action potential. We then averaged all spike-aligned eye positions. Relatively speaking, more eccentric neurons should require larger eye position variation to trigger response variability than less eccentric neurons (e.g. Neurons #6 and #5, respectively, in Fig. 5B, C).

### Shifts in foveal visual RF hotspot locations prior to microsaccades

To analyze the impacts of the second type of fixational eye movement that can take place (microsaccades) on foveal visual representations in the SC, we searched for neurons in our database having sufficient numbers of trials satisfying the following criteria. First, stimulus onset had to occur before microsaccades (within 100 ms), such that we can analyze evoked responses even before a shift in eye position had taken place. Second, there had to be sufficient trials to allow estimating a map of the RF (>10 trials within the RF region). Third, the microsaccades had to be directed opposite the hemifield represented by the SC side that we were recording from, so that we could avoid observing microsaccade-related elevations in firing rate (*56*), which can mask visually-evoked responses by stimulus onset. Fourth, the neurons had to show no microsaccade-related movement burst for these ipsiversive microsaccades. Finally, the neurons had to have RF hotspot locations <2 deg in eccentricity. We then reconstructed visual RF maps without microsaccades (as in all of our analyses) or when stimuli appeared in pre-microsaccadic intervals. Since these intervals are associated with perceptual mislocalizations in the fovea (*17*), we analyzed whether RF hotspot locations were shifted by amounts larger than predicted by simple shifts in eye position caused by the microsaccades themselves. Because all of our calibrations in eye position were done for microsaccade-free fixation (see *Visual and motor-related RF hotspot and area calculations* above), we also confirmed that eye position shifts caused by microsaccades in the present analysis were insufficient to explain our results (e.g. Fig. 6B). We included 24 neurons in this analysis; on average, we had 22 +/- 15.3 s.d. trials for each neuron with microsaccades.

### Estimating a new SC surface topography map from (*37*)

The universally accepted map of SC topography (*37*) was based on peripheral measurements of electrically-evoked saccades from (*65*). We used a similar approach to (*65*), but now densely mapping the most superficial SC laminae in monkeys R and F and with visual stimuli as opposed to electrical microstimulation of the deeper saccade-related layers. We also made sure to measure both peripheral and foveal sites.

We mapped the left SC of monkey F using 500 μm resolution, and using a penetration angle tilted 45 deg posterior of vertical. In monkey R, we mapped the right SC surface topography using 250 μm resolution and vertical penetrations. During the mappings, we also physiologically determined the SC borders, by identifying neighboring sites (Fig. S6A). We performed 91 penetrations, ranging from 0.1 to 49 deg in eccentricity in monkey F, and 124 penetrations, ranging from 0.04 to 12 deg in monkey R. In order to align the SC from the two monkeys, and to obtain a direct comparison to (*37*), we projected monkey R’s recordings into the same 45 deg posterior of vertical coordinate frame (like monkey F). We did this because the original peripheral SC mappings (*65*), based on which the universally accepted SC topography map (*37*) was obtained, were performed using the same approach.

We then ran a rigid body transformation to align recording sites from the two monkeys. In the visual region of overlap between the two monkeys (up to 12 deg eccentricity), we minimized the least-mean-square error between the preferred eccentricity and visual angles of SC sites from the two monkeys. This allowed co-registration of the two SC’s. We validated this co-registration by plotting the non-SC locations that we physiologically measured during the experiments. We confirmed that none of the non-SC sites after co-registration encroached on our aligned SC sites. We also confirmed that there was good agreement in the non-SC sites with known anatomical geometry relative to SC location (e.g. pretectum or cerebellum), and that our map was even consistent with 3-dimensional anatomy of the SC (see *Reconstruction of 3-dimensional SC anatomy* below). The results of the final co-registration are shown in Fig. S6A.

After co-registration, we checked whether foveal sites have organized topography or not, and in a continuous manner with peripheral representation. We identified sites having 0.5 deg preferred visual eccentricity in their visual responses, and we then interpolated between these sites to draw a 0.5-deg iso-eccentricity line. We repeated this procedure for 1, 2, 5, 10, 20, and 40 deg eccentricities (Fig. 3A). We similarly checked iso-direction lines (Fig. 3B).

Finally, to obtain a mathematical fit (*37*), we repeated a procedure similar to that in (*37*). Critically, the key element here was that we had measurements from *both* foveal and peripheral sites, unlike in (*37, 65*). We first rotated the co-registered sites to make the interpolated horizontal meridian (i.e. 0 deg direction line) lie along a horizontal line (e.g. Fig. 3A-D with anatomically-accurate orientation of the horizontal meridian representation versus Fig. 4A with a rotated map to align the horizontal meridian along a horizontal plotting line). We then ran the equations of (*37*), which are:

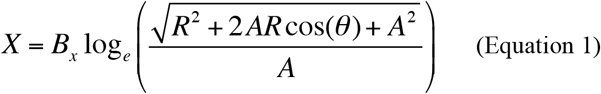

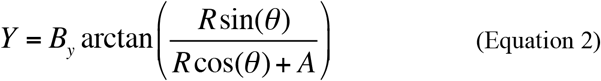

where *X* and *Y* are anatomical distances along the SC axis (*X* being along the axis of the horizontal meridian and *Y* being the axis along the orthogonal axis of directions from horizontal), *R* and *θ* are visual coordinates in eccentricity (*R*) and direction from horizontal (*θ*), and *B*_*x*_, *B*_*y*_, and *A* are model parameters. For a given *R* and *θ* in visual coordinates, *X* and *Y* specify the corresponding locations in SC tissue coordinates. We obtained new model parameters for the above equations by minimizing the least-mean-square error from the equations to our data similarly to the original paper method. Critically, we used all of our measurements for the data fit and not just the iso-eccentricity and iso-direction. We further verified the result by plotting the new parameters on top of our data to visualize the origin of the SC, and we confirmed that the origin was not in the non-SC regions from our dense mappings. The result is shown in Fig. 3C. We also confirmed that our results provide a much better fit than the original model in (*37*) (Fig. 3D). The new model parameters were 1.1 (*B*_*x*_), 1.8 (*B*_*y*_), and 0.9 (*A*).

To estimate the size of the active population of neurons in SC tissue for a given visual stimulus location, transformation of RF shapes from visual coordinates into SC tissue coordinates is needed (*31, 37, 40, 66*). Once this transformation is done, it is trivial to estimate the size of the active population for a given stimulus location: this is achieved by simply checking which neurons in the SC map would be activated by the stimulus (and to what extent according to the stimulus location relative to the RF hotspot location) based on the relative positioning between stimulus location and RF coordinates in neural tissue space (*31, 37, 40, 66*). We thus converted visual RF’s into SC tissue coordinates (Fig. 4A). To do this, we simply plotted firing rate maps as a function of position in the SC tissue space as opposed to visual location. In other words, for every visual location at which we measured RF responses, we converted its eccentricity and direction pair (*R, θ*) from Equations 1 and 2 above into neural tissue coordinates *X* and *Y*.

### Foveal tissue magnification factors

In Fig. 3E, we computed linear foveal magnification factor (mm of neural tissue in the SC map per deg of visual angle in visual space) along the horizontal meridian. We did this for the original universally accepted model of SC topography (*37*) as well as for our new model. We densely sampled the horizontal meridian representation in both models using 0.1 deg visual eccentricity resolution, and we plotted the difference of tissue distance from one sample to the next (appropriately dividing by the step in eccentricity to obtain a correct mm/deg measure).

In Fig. 3F, we wanted to compare the linear foveal magnification factor in the SC to that in the primary visual cortex (V1). However, a direct quantitative comparison of V1 foveal magnification factor to Fig. 3E would have been inappropriate and unfair because the SC is approximately an order of magnitude smaller in size than V1. Therefore, we first scaled the SC size upwards in order to match the size of V1, and we then compared quantitative measures. We scaled the SC upwards based on quantitative measures of V1 size. Specifically, we multiplied SC distance along the horizontal meridian by a scale factor that was based on the ratio of linear distances from 2.5 deg eccentricity to 40 deg eccentricity in both the SC and V1. In the SC, the distance (between 2.5 deg and 40 deg eccentricity lines) from our measurements was 2.8463 mm. In V1, we estimated this distance to be 25.2014 mm based on (*10*). Therefore, we scaled the SC’s horizontal meridian representation by a factor of 8.8541 (equal to 25.2014/2.8463) before comparing SC foveal tissue magnification to that in V1. The comparison was then done by plotting our scaled estimate of SC foveal tissue magnification (if the SC was of approximately the same physical size as V1) to V1 data reported in (*11*) from multiple studies (Fig. 3F).

### Reconstruction of 3-dimensional SC surface anatomy and alignment with normalized postmortem MRI’s

We used dense mappings in monkeys R and F and histological sections to develop a detailed 3-dimensional characterization of macaque SC surface topography. To register the physiological recordings to a common MRI space, we needed an intermediate step to align the anatomical locations of the recordings. In one penetration, we therefore made an electrolytic lesion close to the SC (monkey R; Fig. S6A). We took all histological sections containing SC, and we manually traced the SC border (Fig. S6B). We next interpolated the outline of the SC in 3-dimensions using Free-D (*67*) (Fig. S6B), and we used rigid body transformation to register the reconstructed 3-dimensional SC to a published normalized MRI common space for postmortem macaque monkeys (*68*) (Fig. S6C). We used the cerebral aqueduct as our landmark and for registration with the standard MRI image. After the registration, we used the electrolytic lesion mark from monkey R to register the physiological recordings to the MRI. We also plotted the SC and non-SC regions (as identified physiologically) to confirm that our registration procedure matched with the known anatomical landmarks on the MRI image (Fig. S6D; Movie S1). Finally, we rendered the MRI data with ITK-SNAP (*69*) and plotted our new parameters on the rendered image (Fig. S6E; Movie S2). Data Table S1 contains 3-dimensional coordinates of right SC surface iso-eccentricity and iso-direction lines in our new model of SC topography.

**Figure S1.**
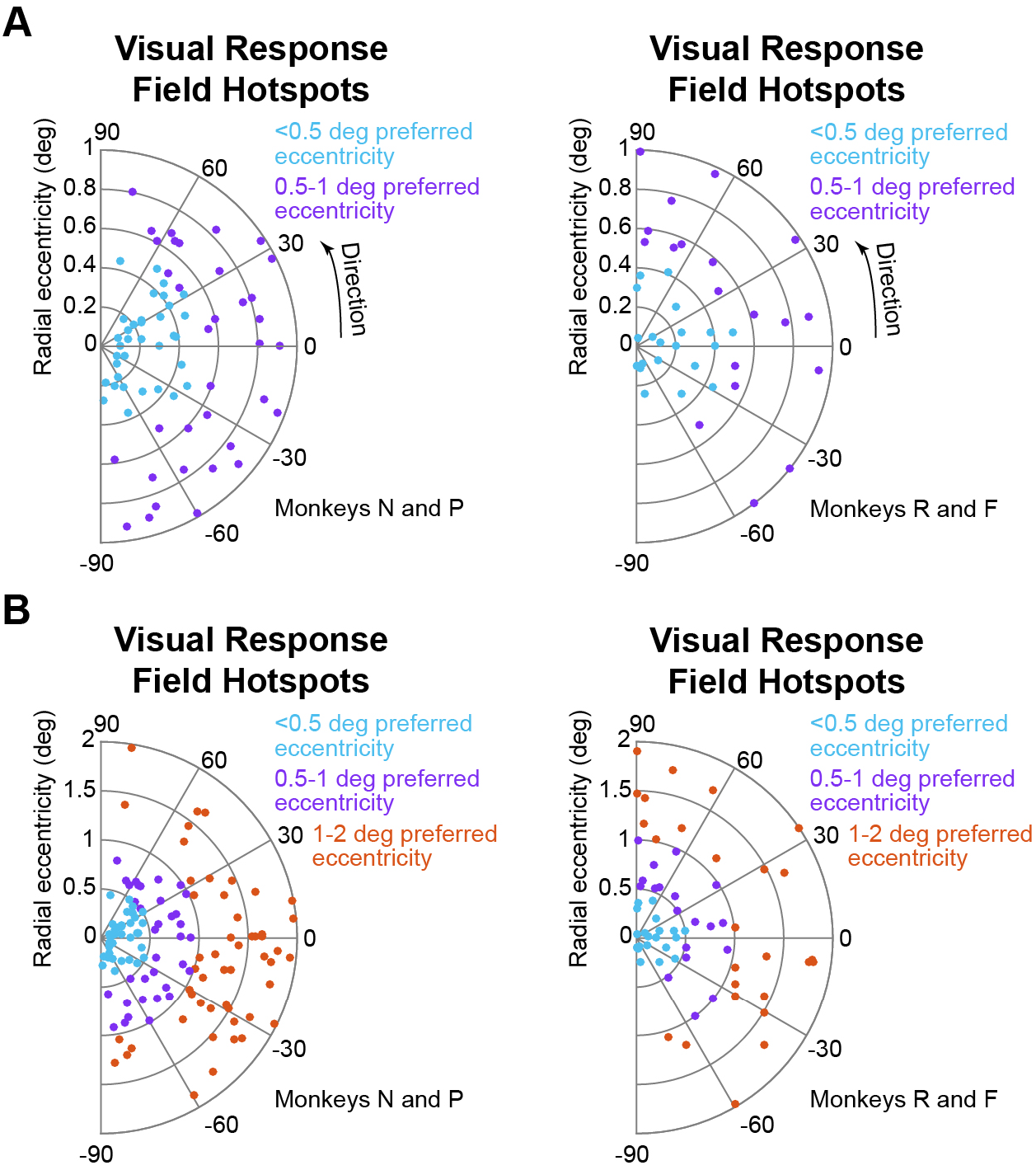
Sampled foveal locations represented by SC neurons in all of our four monkeys. **(A)** The left panel shows preferred RF hotspot locations identified in awake monkeys N and P, and the right panel shows preferred RF hotspot locations identified in anesthetized monkeys R and F. The neurons are color-coded according to their preferred hotspot eccentricity. We sampled a substantial number of neurons representing a region of visual space within the rod-sparse foveola area of the retinal image (<0.5 deg preferred eccentricity). **(B)** The same data as in **A** along with additional neurons (1-2 deg preferred eccentricity), demonstrating approximately uniform sampling of all possible eccentricities <2 deg. More eccentric neurons (>2 deg) are not shown, but they were included in all of our analyses for critical comparisons to foveal SC representations. Note that we recorded in both right and left SC’s in each of monkeys N and P, but we rotated all neurons into one visual hemifield for figure simplicity. All neurons had visual RF hotspots that were located in the contralateral visual hemifield to the recorded SC. For monkeys R and F, we recorded from right SC (representing the left visual hemifield) in monkey R and left SC (representing the right visual hemifield) in monkey F, but we again rotated the data in this figure for presentation purposes.

**Figure S2.**
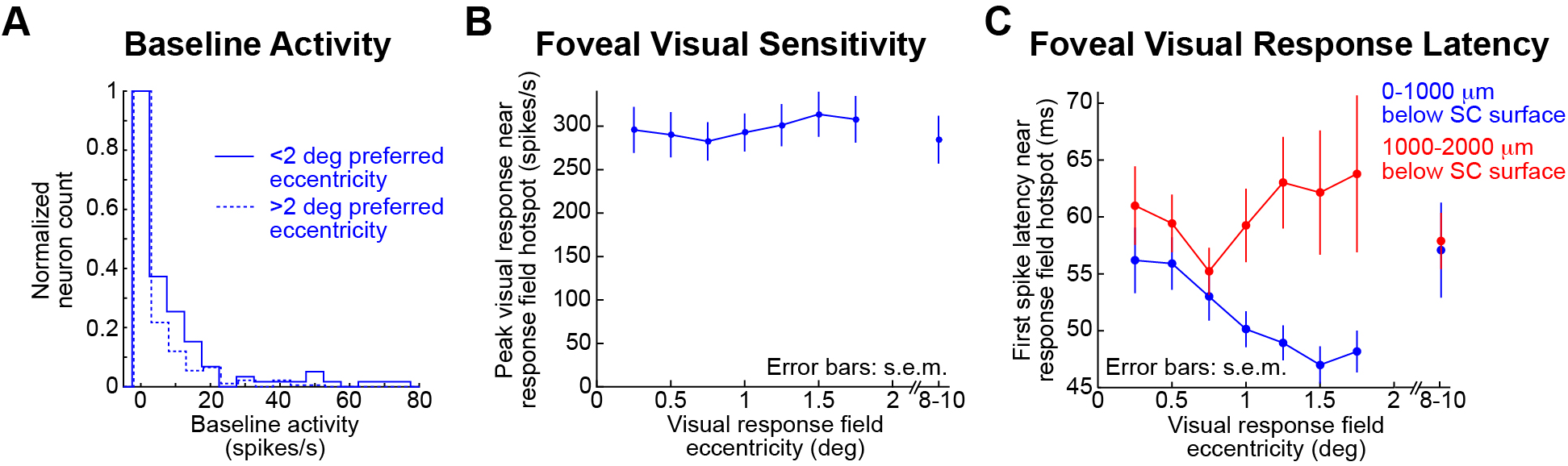
Visual properties of foveal SC neurons. **(A)** Most neurons in the awake animals, whether foveal (solid line) or eccentric (dashed line), had little baseline activity. For each neuron, we measured average firing rate within a 200-ms baseline interval in monkeys N and P (Materials and Methods). **(B)** For all neurons with preferred eccentricity <2 deg (Fig. S1, awake monkeys N and P), we plotted peak visual response near the preferred RF hotspot location (Materials and Methods) as a function of preferred eccentricity. For comparison, we also plotted visual response sensitivity for neurons preferring 8-10 deg eccentricities (again with stimuli appearing at the respective RF hotspots). SC foveal visual sensitivity was as strong as that at 8-10 deg (Ranksum test, p= 0.63753). Also, within the central 2 deg, there was no dependence of visual sensitivity on neuronal preferred eccentricity (1-way ANOVA, F=0.1924, p=0.9014). **(C)** We measured the latency to first stimulus-evoked action potential (spike) for the same locations and neurons as in **B**. In this case, we separated the neurons as being either superficial (<1000 μm below SC surface) or deeper (1000-2000 μm below SC surface) because they showed different results. Superficial neurons (blue), which are retina-recipient (*3-5*), had longer first-spike visual response latencies within the foveola region and decreased their response latency with increasing eccentricity in the rest of the central 2 deg (1-way ANOVA, F=3.4096, p=0.023). This might reflect an impact of rod-sparse foveolar cone input to neurons (whether directly or through cortex) representing <0.5 deg eccentricities. Deeper neurons (red), not receiving direct retinal input, did not show a dependence of first-spike latency on eccentricity in the central 2 deg (1-way ANOVA, F=0.7479, p=0.5288). Note also that superficial neurons responded earlier than deeper neurons in general (Ranksum test, p=0.0040, all eccentricities <2 deg; p=6.375×10^-6^, all eccentricities), consistent with peripheral results (*30*). Error bars denote s.e.m.

**Figure S3.**
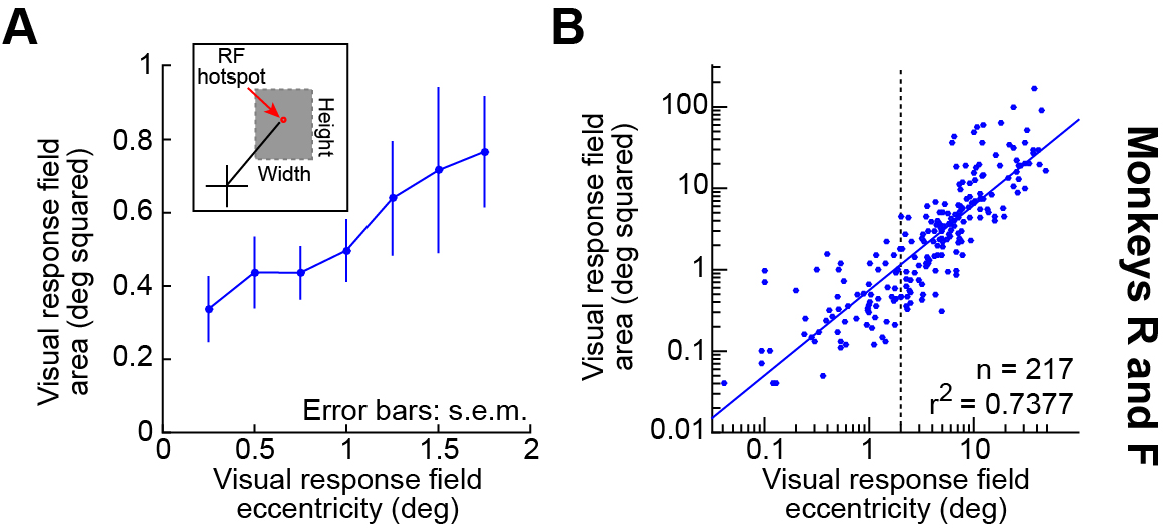
Non-uniform sampling within the SC foveal visual representation even in anesthetized monkeys. **(A)** For monkeys R and F, we performed similar analyses on visual RF area to those presented in Fig. 2A for the awake monkeys. The inset shows how we assessed RF area in these anesthetized cases (Materials and Methods). Error bars denote s.e.m. (p=0.05, 1-way ANOVA as a function of eccentricity, F=2.75, n=66). **(B)** Data from the central 2 deg are now shown along with the rest of the neurons that we sampled. Each dot represents a single neuron (as in Fig. 2A), and we now plot data on a logarithmic scale because of the large range of eccentricities and areas that we measured. The blue line shows a linear regression fit of all data points (on the logarithmically transformed scales). Like with the awake monkeys (Fig. 2A), the SC foveal visual representation formed a continuum, in terms of spatial sampling resolution, with the peripheral representation, showing a supra-linear growth in RF area as a function of increasing eccentricity. The dashed vertical line delineates the 2 deg eccentricity line. Foveal neurons (<2 deg preferred eccentricity) clearly showed non-uniform sampling resolution, like in the peripheral representation, and also like in the awake animals (Fig. 2A). Note that we did not quantitatively compare RF areas between awake and anesthetized animals due to several experimental constraints (Materials and Methods).

**Figure S4.**
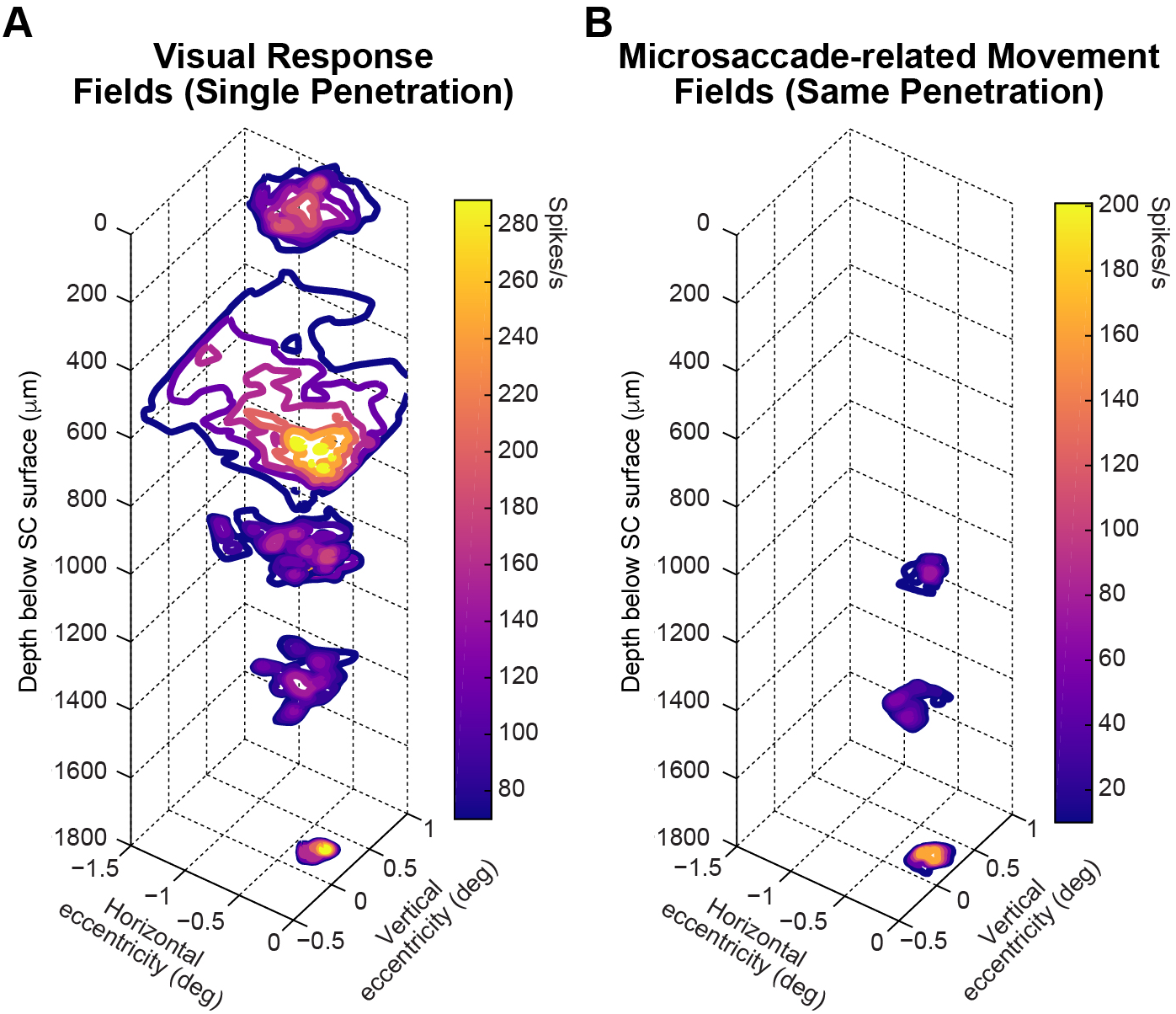
Consistent vertical organization of visual and visual-motor neurons within the SC foveal representation. **(A)** Example visual RF maps from 5 different neurons isolated within the same recording session in an awake monkey, as the recording electrode was advanced deeper and deeper into the SC (Materials and Methods). Each plane along the z-axis plots a pseudocolor surface describing peak visual response after stimulus onset (color-coded) as a function of horizontal and vertical target eccentricity (x- and y-axes). The z-axis plane at which each surface is drawn indicates the depth at which the neuron was encountered relative to SC surface. As can be seen, neurons within a single “depth column” of the SC had similar preferred eccentricities, consistent with peripheral SC representations. **(B)** The three deepest neurons from **A** also had microsaccade-related movement RF’s (Materials and Methods) (*15, 56*). Thus, the most superficial two neurons were purely visual neurons (no movement-related discharge), whereas the deeper three neurons were visual-motor neurons. This ordering of visual and visual-motor neurons is consistent with peripheral SC depth-related organization (*70*).

**Figure S5.**
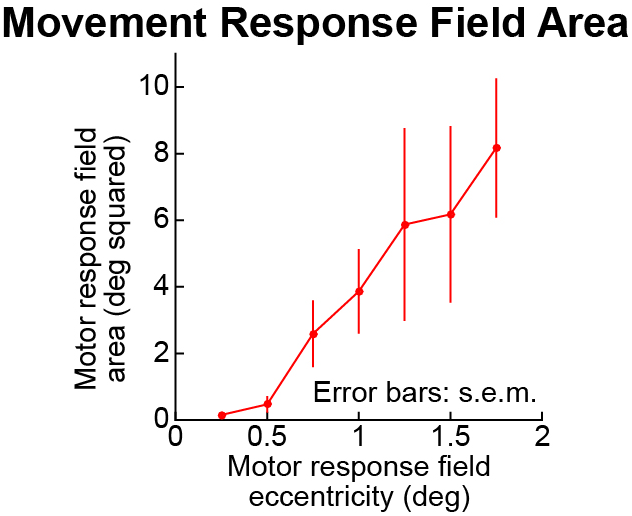
Non-uniform spatial sampling within the SC representation of foveal space even for movement-related RF’s. For the awake monkeys N and P, we measured microsaccade- and saccade-related responses (Materials and Methods), and we performed the same RF area analyses on these responses. Within the foveal representation, we observed similar non-uniform sampling resolution, in terms of RF area, that we observed for visual RF’s (e.g. compare to Fig. 2A and Fig. S3) (p=1.260×10^-3^, 1-way ANOVA as a function of eccentricity, F=6.05, n=58). Note that microsaccade-related movement RF’s in the fovea (<1 deg) were substantially smaller than visual RF’s in the same animals (Ranksum test, p=2.180×10^-5^) (*15, 56*). Error bars denote s.e.m.

**Figure S6.**
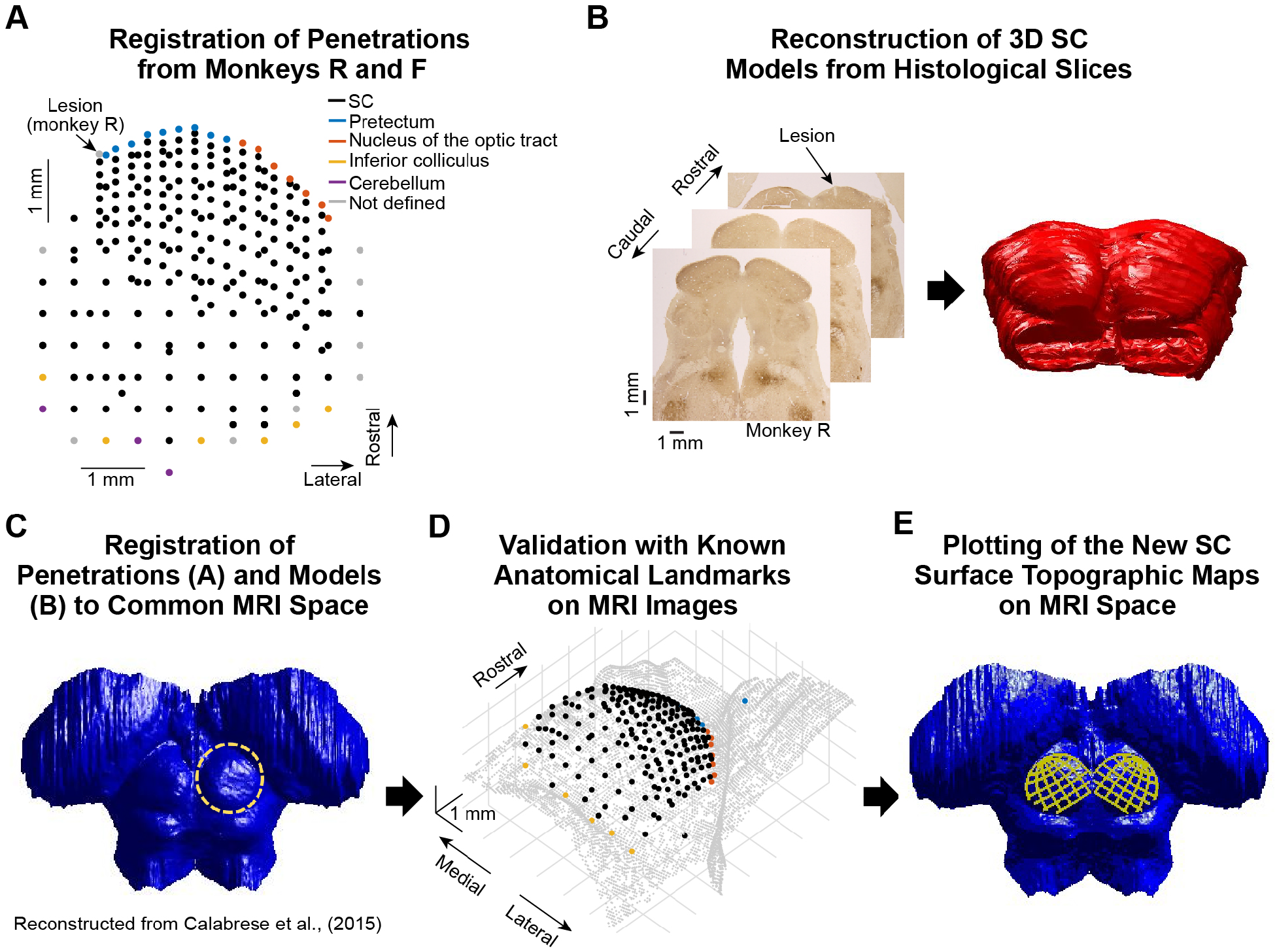
Reconstruction of 3-dimensional SC surface anatomy and topography. **(A)** We co-registered dense mappings from monkeys R and F (black dots; Materials and Methods), and then confirmed that physiologically-identified non-SC sites were at appropriate anatomical locations relative to the SC (Materials and Methods). Each dot represents a single electrode penetration reaching either the most superficial SC laminae (black) or non-SC sites (colors). The colored sites are SC-neighboring structures (legend). **(B)** Histological sections were aligned using an electrolytic lesion and then manually traced to generate a 3-dimensional structure (Materials and Methods). The lesion was placed 100 μm rostro-medially to the last visual RF in the SC. **(C)** The data were then aligned to a standardized postmortem macaque MRI sequence (*68*) (Materials and Methods). **(D)** This allowed us to map our electrode penetrations from **A** to 3-dimensional structure, and to confirm the anatomical landmarks based on SC-neighboring structures (see Movie S1). **(E)** We used our new SC surface topography model (Figs. 3-4) to describe 3-dimensional SC surface topography on a standard macaque MRI (see Movie S2). The left SC topography shown is just a mirror image of the right SC topography that we generated from our data. Data Table S1 lists all 3-dimensional coordinates for the shown iso-eccentricity and iso-direction lines in the right SC as a reference for future uses of the 3-dimensional model.

**Figure S7.**
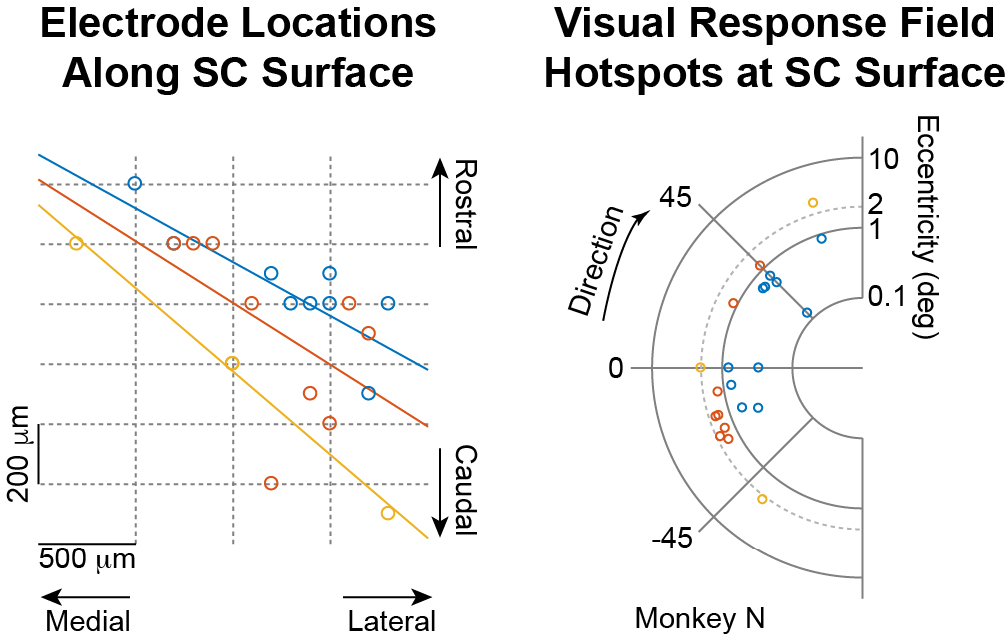
Orderly topographic organization within SC foveal visual representation even in awake monkeys. We took a contiguous series of 22 back-to-back awake-monkey recording experiments, in which we systematically sampled within the rostral region of a single SC (the right SC) of a single monkey (monkey N). In the left panel, we plotted penetration locations within the recording chamber (the chamber was placed on the midline and tilted 35 deg posterior of vertical in this animal; Materials and Methods), and we color-coded the penetration locations according to the preferred visual eccentricities encountered in the most superficial SC laminae. The preferred eccentricities are plotted in visual coordinates in the right panel (using a logarithmic eccentricity scale). The colored lines in the left panel are linear regression lines for each color coding of penetration location, demonstrating orderly clustering of iso-eccentricity sites, and with an anatomical orientation (relative to the rostral-caudal axis) similar to that seen in the anesthetized animals (Fig. 3A-D). We also observed that medial sites represented upward directions from horizontal and lateral sites represented downward directions. Note that a direct comparison of the left panel to Fig. 3 is not appropriate because of the different penetration angles in the two cases.

**Figure S8.**
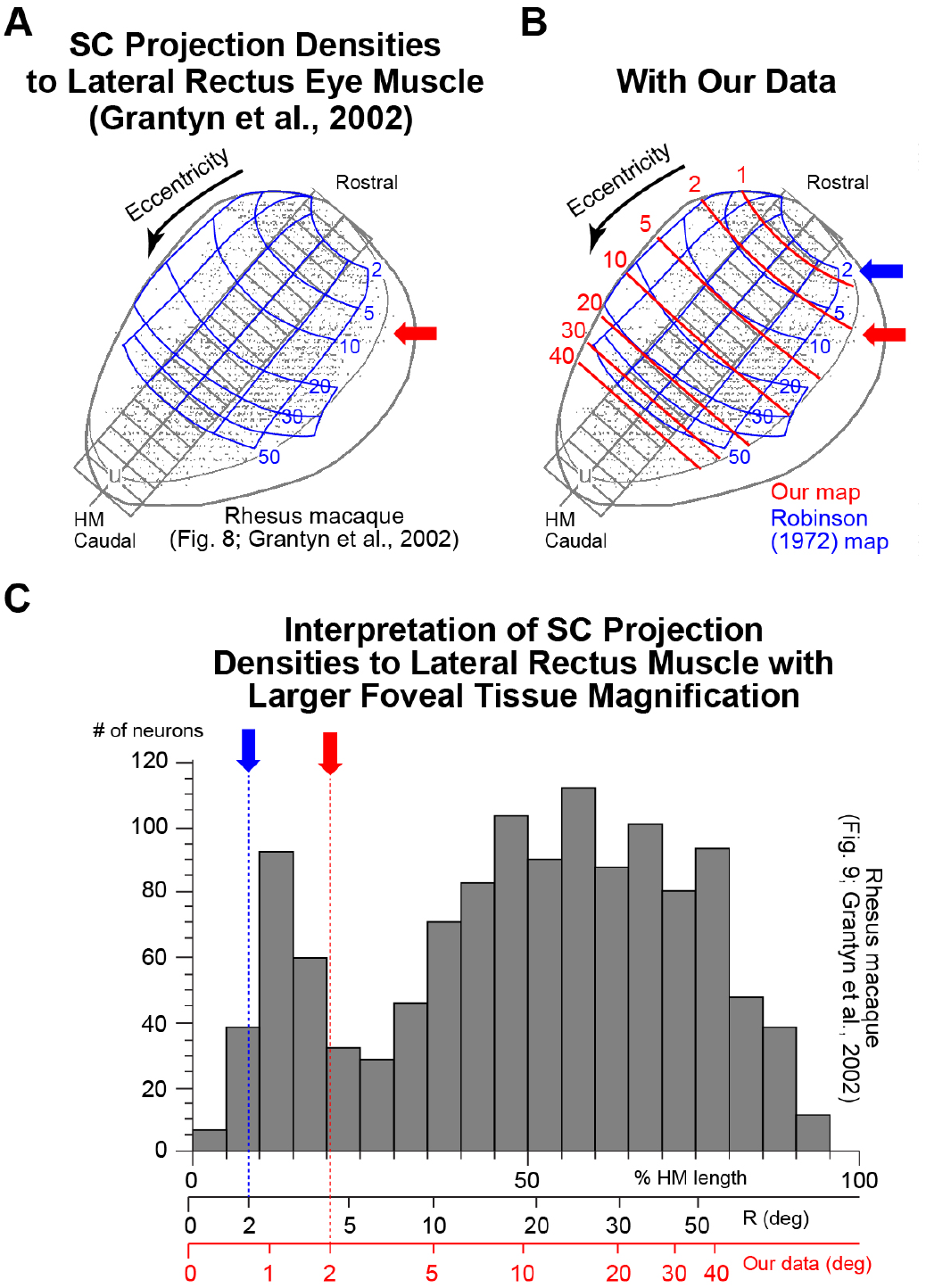
Recasting interpretations of SC efferent projections to the oculomotor periphery. **(A)** Besides comparisons to primary visual cortex (V1), a potential source of afferent input to the SC (Fig. 3F), we also explored the implications of larger foveal magnification on interpreting efferent SC projections. The figure, adapted with permission from (*38*), shows SC projection densities (gray dots) to the lateral rectus eye muscle (driving horizontal eye movements). Blue is Grantyn et al.’s (*38*) estimate of topography, based on (*65*), and the diagonal grid is their binning of tissue for the measurements in **C**. HM means horizontal meridian. The red arrow shows the 2-deg eccentricity line according to our more complete measurements of SC topography (Figs. 3-4, S6, S7), if our map was aligned at the origin with the Robinson map alignment (as in Fig. 3D); this demonstrates how the interpretation of raw cell densities (gray dots) can be grossly misestimated. **(B)** This is better illustrated when our map is superimposed in red, again when the origin was aligned to the same origin of the blue map. **(C)** Histograms of the cell densities from **A**, **B** along the horizontal meridian. With the original map placement (black horizontal axis of eccentricity), it may appear that the central 5 deg of SC representation are a distinct zone because of the bimodal distribution dip near 5 deg in the black horizontal axis. However, with our more complete mapping of SC topography, the distinct zone transition in cell densities may delineate a much smaller eccentricity (red arrow).

**Figure S9.**
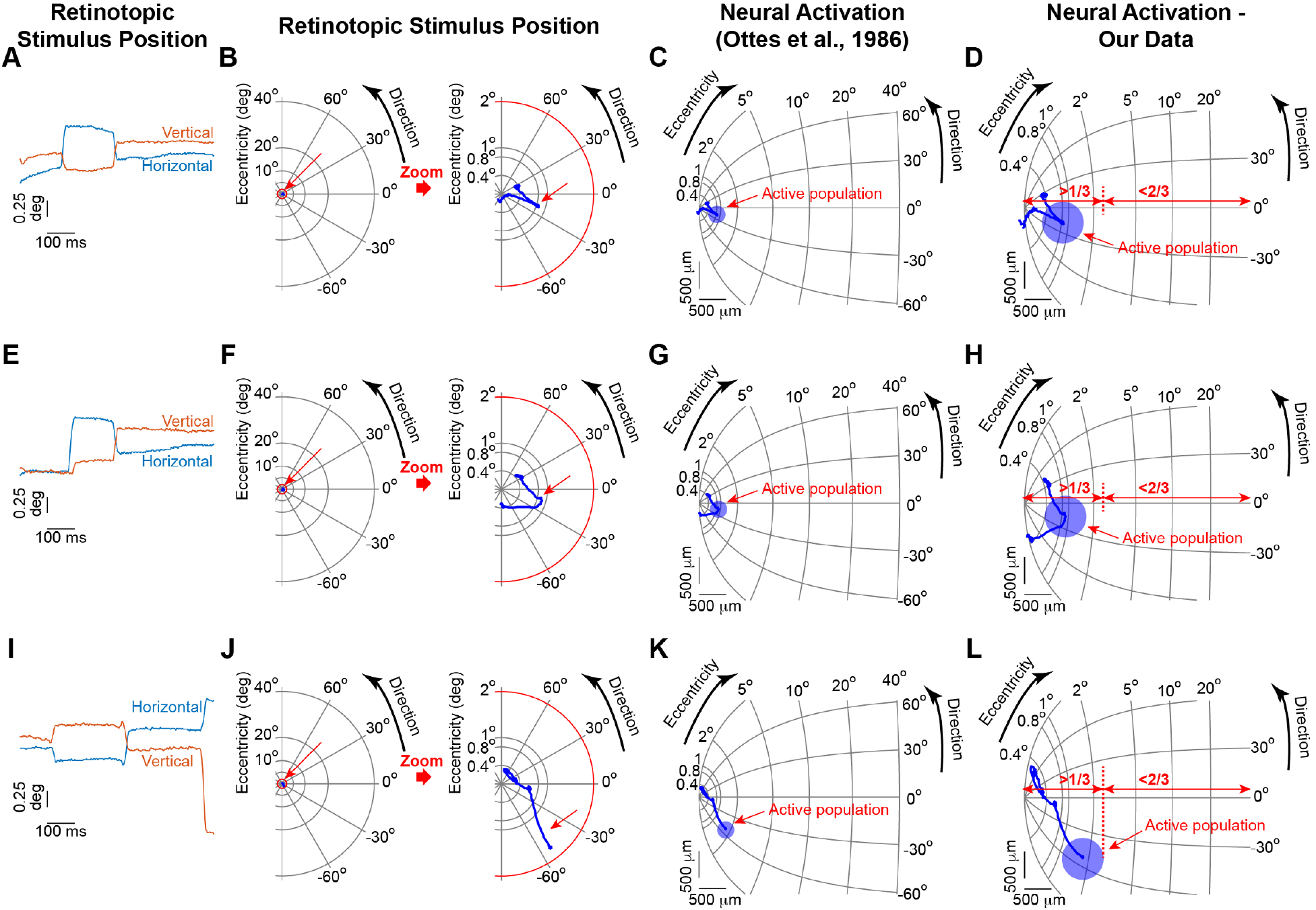
SC activation when tiny fixational eye movements shift retinal images. The combination of foveal magnification (Figs. 3, 4) and eye movements (Figs. 5, 6) means that even small foveal stimuli can cause significant activation waves. The figure shows how the retinal image of a small foveal spot gets moved around by fixational eye movements and what this means in neural tissue coordinates. **(A)** Retinotopic stimulus position in one example trial of a small stationary spot (e.g. Fig. 5A). **(B)** The left panel shows the motion trajectory (blue) from **A** in retinotopic visual field coordinates. The spot was initially place at 0.4 deg before being shifted around by fixational eye movements, and the small red circle delineates 2 deg. Relative to the entire visual field (left panel), the motion trajectory (blue) was barely visible. The right panel magnifies the central 2 deg to highlight the small motion. **(C)** The motion of the spot on SC tissue as per the classic model of (*37*). The blue circle demonstrates an estimate of the active population (e.g. Fig. 4) at a specific time during the trial. **(D)** A more accurate depiction of stimulus trajectory from the same trial based on our dense mapping of SC foveal topography (Figs. 3, 4). A simple foveal stimulus can activate up to ∼1 mm of SC tissue due to small fixational eye movements. **(E-H)** Same for another example trial. **(I-L)** Same for another example trial. Note how the edge of the active population (red dashed vertical line) can reach a distance greater than 1/4-1/3 of the entire SC rostral-caudal extent. Also note that in real neural dynamics, there can be even longer range interactions than simply stimulus position (*42, 43*). SC maps were rotated as in Fig. 4 such that the eccentricity dimension was on the horizontal dimension.

**Movie S1**

Results of Fig. S6D after aligning our penetration locations with anatomical locations of in the SC. The color coding for non-SC sites is the same as that in Fig. S6A.

**Movie S2**

Results of Fig. S6E. Iso-eccentricity and iso-direction lines when project onto the 3-dimensional anatomical SC structure. Data Table S1 lists all 3-dimensional locations in this movie for future uses of our revised view of SC topography. The left SC iso-eccentricity and iso-direction lines were reflected from measurements of the right SC.

**Data Table S1**

A Microsoft Excel file (related to Fig. S6 and Movie S2) containing tables of x, y, and z coordinates for the right SC’s anatomical coordinates along with individual iso-eccentricity and iso-direction lines. The file contains a table for iso-eccentricity lines and a table for iso-direction lines. The file also contains additional tables with coordinates for the 3-dimensional anatomical SC surface (for generating Movie S2), as a resource for future uses of our SC surface topography. All measurements are in millimeters (mm) relative to the anterior commissure, as explained in the file (*68*).

